# Coordination-dynamics invariants from simultaneous vagus nerve electroneurogram and ECG: a Python pipeline applied to neonatal piglet endotoxaemia

**DOI:** 10.64898/2026.06.01.728499

**Authors:** Martin G. Frasch, Mike Last, Patrick Burns, Gilles Fecteau, Andre Desrochers

## Abstract

HRV complexity metrics track the systemic cytokine response across mammalian inflammation models, but the dynamical mechanism translating molecular inflammation into HRV structure has remained unresolved, and no existing tool jointly extracts coordination-dynamics signatures from simultaneous vagus nerve electroneurogram (VENG) and ECG. We describe a modular open-source Python pipeline that (i) detects spectrally distinct latent states in 20 kHz whole-trunk VENG using a symmetric-Kullback–Leibler scan-statistic changepoint detector ported from earthquake seismology and verified against its R progenitor; (ii) places the inter-changepoint-interval series in the Haken–Kelso–Bunz coupled-oscillator framework, computes the Kuramoto order parameter *R*, fits the bistable potential *V*(*φ*) = −*a* cos *φ* − *b* cos 2*φ*, and maps each timepoint into the Arnold-tongue (Ω, *K*) plane; contrasts compound-action-potential band power between successive latent states; and (iii) couples the VENG analysis to a CIMVA HRV pipeline driven by a 5-algorithm Pro-MAC R-peak ensemble. Applied to a previously published two-animal neonatal piglet endotoxaemia cohort (Castel et al., 2020, 2024), the pipeline recovers known features of the (iv) preparation and surfaces three case-series observations that we frame as hypotheses for prospective replication: (1) a 6–8 s VENG state-switching period coincident with the HRV LF/HF spectral boundary; (2) qualitatively divergent coordination-dynamics trajectories under LPS versus LPS + VNS (Kuramoto *R* collapses from 0.69 to 0.12 without VNS, undergoes biphasic recovery to 0.99 with VNS, the two trajectories crossing at 60–75 min); (3) a regime-dependent empirical coupling between *R* and the HRV embedding scaling exponent eScalE (*R* · eScalE ≈ 0.33 during moderate challenge, doubling to ≈0.80 during VNS overshoot). Pre-specified null comparisons (Poisson and bootstrap surrogates) demonstrate that *R* and the critical-slowing-down indicators carry information beyond their most obvious null alternatives. With *N* = 2 animals no inferential statistics are possible, so these findings are reported as hypothesis-generating; the manuscript closes with a pre-registered prospective protocol designed specifically to falsify them.

**Graphical Abstract:** 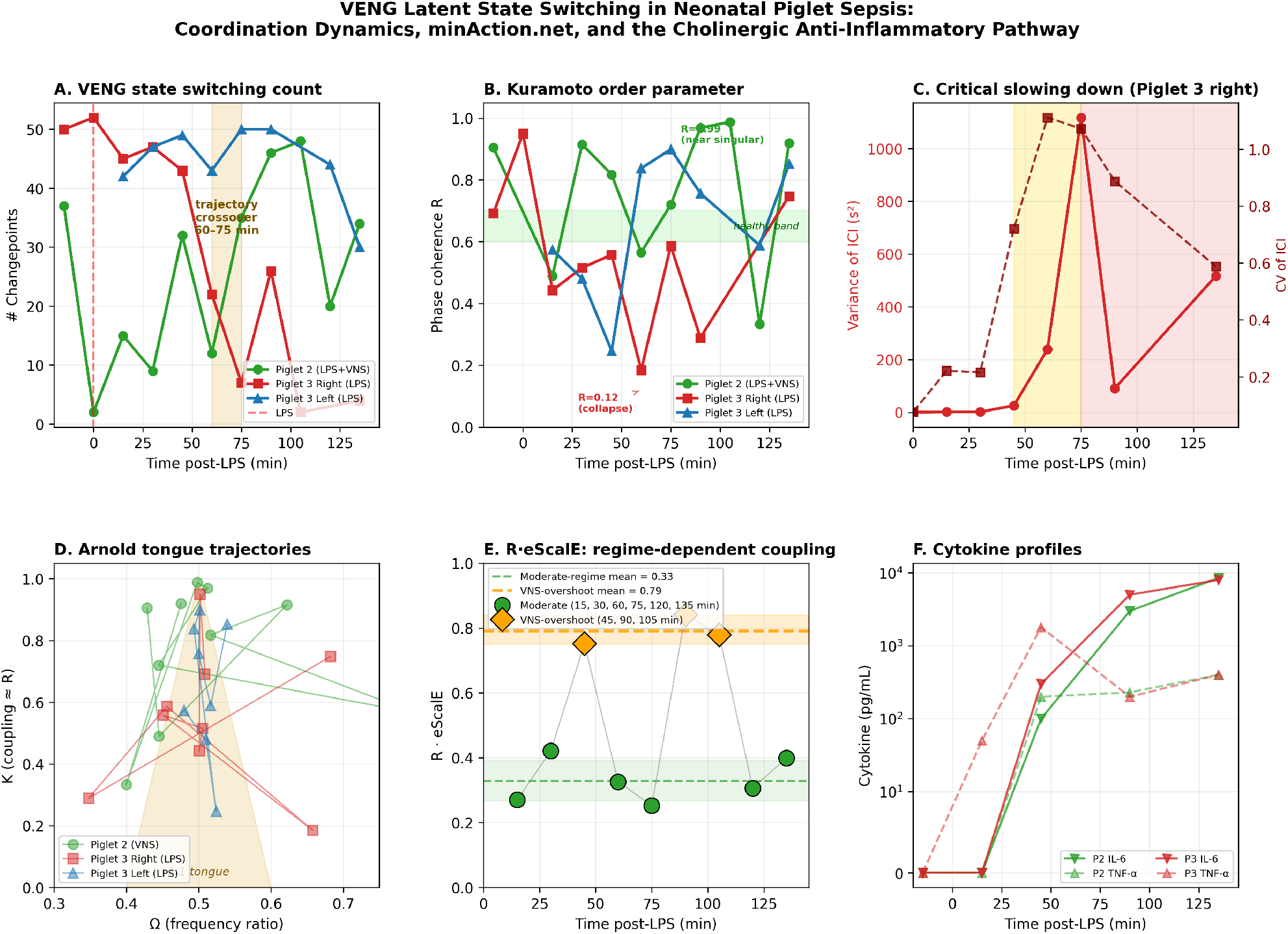

**Key Points:**

- We present an open-source Python pipeline that extracts coordination-dynamics invariants (Kuramoto phase coherence, Haken–Kelso–Bunz potential parameters, Arnold-tongue position, critical-slowing-down indicators) from simultaneously recorded vagus nerve electroneurogram (VENG) and ECG, starting from a spectral-KL scan-statistic changepoint detector ported from earthquake seismology.
- Applied to a previously published two-animal cohort (Castel et al., 2024), the pipeline reveals that VENG alternates between two spectrally distinct latent states at a baseline period of 6–8 s that coincides with the HRV low-frequency/high-frequency spectral boundary (∼0.15 Hz). We propose the VENG latent-state oscillator as a *candidate* mechanistic generator of vagal LF HRV; this is a hypothesis motivated by the timescale match and is not established by the present data.
- In this cohort, LPS endotoxaemia and vagus nerve stimulation drove qualitatively divergent trajectories in the coordination-dynamics phase space (Kuramoto *R*: 0.69 → 0.12 without VNS; biphasic to *R* → 0.99 with VNS at the IL-6/IL-8 cytokine peak). Because the design provides one animal per arm, these are case-series observations and not a between-arm statistical comparison.
- Inter-changepoint-interval variance rose >100-fold roughly 15 min before overt state-switching collapse in the unmitigated animal. Bootstrap nulls drawn from the same animal’s pre-collapse ICI pool reject the same-distribution alternative (§2.6), motivating a critical-slowing-down early-warning hypothesis that requires prospective replication before any clinical use is warranted.
- The pipeline surfaces a regime-dependent empirical coupling between VENG phase coherence (*R*) and HRV embedding scaling exponent (eScalE) in the VNS-treated animal: during moderate challenge *R* · eScalE ≈ 0.33 (CV ≈ 0.20), during VNS-enhanced overshoot (45, 90, 105 min) the product doubles to ≈0.80. The product is a post-hoc empirical construct, not a derived conservation law; we propose it as a falsification target for prospective cohorts (§6).

## 1 Introduction

The cholinergic anti-inflammatory pathway (Tracey, 2002) established the vagus nerve as a regulator of systemic cytokine production, and vagus nerve stimulation (VNS) has since become a central paradigm of bioelectronic medicine for controlling inflammation across diverse animal models and early clinical studies (Kwan et al., 2016). Cytokine-specific neurograms recorded directly from the vagus (Steinberg et al., 2016) demonstrated that the molecular inflammatory state can in principle be read from the nerve’s electrical activity. In parallel, we and others have shown that the same information is recoverable non-invasively from heart rate variability (HRV): a specific combination of complexity features (eScalE, sgridTau, AsymI, Multifractal_c1), combined via principal-component analysis into the CIMVA inflammatory index, tracks the IL-6 temporal profile in near-term fetal sheep (Durosier et al., 2015; Herry et al., 2016) and—at a three-orders-of-magnitude higher LPS dose—in neonatal piglets (Castel et al., 2020, 2024). The pivotal cross-check in that piglet work was that the same metrics applied to the VENG signal itself produced a VENG-derived inflammatory index that tracked the cytokine profile in close agreement with the HRV-derived one, indicating that the mathematical signature exists already in the nerve.

Tissue-level work from our group has further shown that the vagus regulates *immunometabolic* homeostasis at the level of the fetal ileum (Cao et al., 2024), that microglial and macrophage plasticity and regional cerebral blood flow in the prenatal brain and gut respond to vagal manipulation (Wakefield et al., 2025), and that the methodological substrate for chronic cervical VENG recording and stimulation in large-animal preparations is now fully described (Castel et al., 2021). What these complementary lines of work have *not* yet produced is an integrated analytical tool capable of extracting, from simultaneously recorded VENG + ECG, the joint dynamical-systems structure that allows such multi-scale translation to be tested as a coordination-dynamics mechanism rather than as a correlation.

The conceptual motivation for seeking such a tool comes from our recent causal framework (Frasch, 2026): we argued there that multi-scale physiological mechanisms are best sought not in horizontal, scale-specific chains but in *vertically organizing principles*—scale-invariant action-like functionals whose minima are the system’s stable operating points and whose deformations encode perturbations. That framework makes three concrete empirical predictions for any physiological subsystem that is vertically organized: (i) it should exhibit coordination-dynamics signatures (mode-locking, bifurcations, critical slowing); (ii) perturbations crossing a bifurcation threshold should produce *qualitatively* different trajectories rather than graded linear responses; and (iii) there should exist approximate conserved quantities linking scale-specific state variables. The pipeline described here was built specifically to test these predictions on the cholinergic anti-inflammatory pathway using an existing piglet endotoxaemia dataset.

The contribution of this *Techniques for Physiology* article is therefore threefold. Methodologically, we describe and release a modular Python pipeline (veng_minaction) that takes raw VENG and ECG as input and returns a full coordination-dynamics characterisation together with a joint VENG–HRV empirical-coupling scan. Physiologically, we demonstrate on a previously published two-animal neonatal piglet cohort that the pipeline recovers all known qualitative features of the preparation while surfacing three new case-series observations—the 7-s state-switching oscillator, the qualitatively divergent within-animal effect of LPS and VNS on coordination-dynamics parameters, and a regime-dependent *R* · eScalE coupling—that were not accessible with any prior tool. We are explicit throughout that with *N* = 2 animals and asymmetric data quality these are hypothesis-generating observations, not inferentially established findings. Translationally, we specify a pre-registered prospective protocol designed to falsify these observations in a properly powered cohort.

## 2 The Technique

The pipeline is implemented as a set of loosely coupled Python modules under veng_analysis/. All parameters are exposed in a single YAML configuration file, and every analytical step returns serialisable artefacts (NumPy arrays, pandas DataFrames, Matplotlib figures) to support reproducibility and independent reanalysis.

### 2.1 Input format and preprocessing

The pipeline accepts EDF/EDF+ and Spike2 TXT exports. EDF files lacking valid physical calibration (blank physical minimum/maximum)—a common real-world problem in chronic large-animal recordings—are loaded through a fallback raw-digital reader that returns int16 samples and flags the channel as uncalibrated; downstream modules then refuse absolute cross-animal amplitude comparisons but accept within-animal temporal analyses. Spike2 TXT exports are parsed by a multi-channel extractor that identifies VENG, ECG, and BP channels by heuristic labelling and returns per-channel DataFrames with provenance metadata.

### 2.2 Spectral-KL scan-statistic changepoint detection

The core detector is a Python port (verified numerically against the original R code on matched baseline recordings) of a spectral changepoint scan statistic originally developed for seismic signals. The detector scans the signal in steps of *skip* samples; at each candidate centre *t* it computes smoothed periodograms *P*_*L*_ and *P*_*R*_ on left and right half-windows of length *bandwidth* and reports the symmetric Kullback–Leibler score

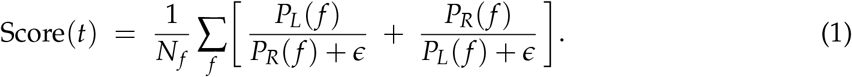

A greedy local-maxima suppression with user-defined score threshold and suppression radius returns the final changepoint set. Default parameters for 20 kHz VENG are *bandwidth* = 65 536 samples (≈3.28 s), *skip* = 2 000 samples (100 ms resolution), smoother = 101 taps, frequency partition at 800 Hz, score threshold = 2.05, suppression radius = 33 score indices; all are exposed in the YAML file and all results in §3 are reproducible with this single parameter set. The detector is agnostic to the absolute scale of the signal, which is the key property that allows it to operate on the uncalibrated Piglet 3 recordings.

### 2.3 Coordination-dynamics wrapper

Given the detected changepoints, the pipeline treats the sequence of inter-changepoint intervals (ICIs) as successive periods of a state-switching oscillator and derives four families of coordination-dynamics descriptors. The “two latent states” referred to throughout are the alternating segments *between* successive changepoints: each detected changepoint is the boundary between two segments whose smoothed periodograms are maximally divergent in symmetric Kullback–Leibler distance, so neighbouring segments differ in spectral content by construction. Even-indexed and odd-indexed segments are then treated as the two states for the CAP test (§2.4) and the Arnold-tongue Ω definition; we make no a priori commitment to which spectral signature corresponds to which physiological mode, only that the alternation pattern is statistically distinct from a stationary process.

#### Phase-coherence index *R* (analogous to a Kuramoto order parameter)

For each successive ICI value we assign an instantaneous phase *θ*_*i*_ = 2*π* ICI_*i*_/⟨ICI⟩ relative to the empirical mean ICI (the expected switching period), and compute *R* = |⟨exp(*iθ*_*i*_)⟩|. So defined, *R* measures the *phase concentration of the ICI distribution around the mean*: *R* = 1 corresponds to a perfectly metronomic two-state oscillator (all ICIs equal), *R* → 0 corresponds to ICIs uniformly distributed in phase (e.g. Poisson-like switching), and *R* ≈ 0.6–0.7 is the empirically observed healthy-range band in this preparation (sub-second jitter on a ∼6.5–8 s switching period). We adopt the Kuramoto-style notation (Kuramoto, 1984) by analogy because the same algebraic form is used in the original coupled-oscillator setting, but emphasise that this is *not* a Kuramoto model in the population-synchrony sense—there is one observed time series, not an ensemble of phase oscillators. *R* is invariant to the absolute switching timescale and depends only on relative ICI dispersion, which is the property that licences cross-animal comparison.

For Gaussian ICIs with coefficient of variation CV ≪ 1 a closed-form expectation *R* ≈ exp[−(2*π* · CV)^2^/2] holds, so *R* is monotonically related to CV in that limit. Whether *R* might therefore be redundant with CV is addressed empirically in §2.6: across the full observed range the rank correlation *ρ*(*R*, CV) = −0.33, so *R* is *not* simply a transformed CV—it tracks non-Gaussian structure of the ICI distribution that CV alone does not register.

#### HKB bistable potential

We fit the Haken–Kelso–Bunz potential *V*(*φ*) = −*a* cos *φ* − *b* cos 2*φ* (Haken et al., 1985) to the ICI phase distribution and report the control parameter k = b/a. The bistable-regime threshold is *k* = 0.25: at *k >* 0.25 the potential has two stable wells and the two-state oscillator is dynamically robust; at *k <* 0.25 the second well disappears and one state dominates. The barrier height Δ*V* between wells is reported as an additional resilience measure (deeper wells ⇒ harder to perturb out of the bistable regime). *k* and *R* are independent estimators of the same underlying coupling strength but are computed from different features of the ICI series—phase-difference distribution shape vs. event-wise phase regularity—so their concordance is a methodological cross-check.

#### Arnold-tongue mapping

Each recording segment is mapped into the (Ω, *K*) plane, where Ω is the relative duty-cycle weight between the two latent states (defined operationally as Ω = ⟨ICI_even_⟩/[⟨ICI_even_⟩ + ⟨ICI_odd_⟩], where the ICI sequence is split into even-indexed and odd-indexed alternations) and *K* ≈ *R* is used as the coupling coordinate. Ω = 0.5 corresponds to symmetric alternation between the two states (each holds the same average dwell time); Ω → 0 or Ω → 1 corresponds to one state dominating. The 1:1 mode-locking tongue is centred at Ω = 0.5; recordings inside the tongue display stable two-state alternation and recordings outside it display drift or desynchronised switching. Healthy baseline trajectories sit near Ω ≈ 0.5 with *K* in the 0.6–0.9 range; perturbations are evaluated by whether they move the system further into the tongue (deeper *K* at fixed Ω) or out of it (combined drift in Ω and collapse in *K*).

#### Critical-slowing-down indicators

The variance and coefficient of variation of ICIs are computed in sliding windows of user-selected width. The CV of ICIs is the dimensionless companion of *R*: in the healthy band *R* ≈ 0.6–0.7 corresponds to CV ≈ 0.3–0.4, while a CV approaching or exceeding 1.0 marks Poisson-like switching and is the operational signature of an oscillator at or past the bifurcation. A >2-fold rise in CV in successive windows is flagged as a bifurcation precursor in the spirit of Scheffer et al. (2009); the early-warning window is the time interval between the precursor flag and the overt collapse of the changepoint count.

### 2.4 CAP even/odd contrast

A compound-action-potential (CAP) analysis wrapper adapted from Steinberg et al. (2016) high-pass filters the VENG at >160 Hz, detects CAPs at a 3× local-RMS threshold within a 0.5– 5 ms duration window, and computes per-segment band power in three bands (low ≤ 1 kHz, medium 1–2 kHz, high 2–10 kHz). Successive inter-changepoint segments are then labelled even/odd and compared per-band with a Mann–Whitney *U* test. The expected null is that the two latent states carry the same band-power distribution—i.e. the changepoint detector has merely segmented a stationary process. A significant even/odd difference in the 2–10 kHz band is therefore interpreted as evidence that the two latent states are *functionally distinct* neural states rather than statistical artefacts of the segmentation. The test does not identify which neural mode either state corresponds to; specific physiological labels (e.g. afferent-sensing vs. efferent-output phases of a vagal duty cycle) are speculative interpretations discussed in §4 and are not established by the band-power comparison alone. Conversely, the absence of a significant band-power difference would indicate that the two states are functionally equivalent and that the changepoint structure is purely statistical—a falsifiable prediction the test is designed to distinguish.

### 2.5 Joint VENG–ECG empirical-coupling scan

The final module operates on *simultaneously* recorded VENG and ECG. ECG is processed with a 5-algorithm ProMAC R-peak ensemble (Pan–Tompkins, Hamilton, Christov, Engelse–Zeelenberg, and neurokit2 default) with per-beat signal-quality index (SQI) gating; the SQI is the agreement fraction across the five detectors, and beats below threshold are rejected before downstream analysis. The resulting RR interval series is passed through a reimplementation of the CIMVA metrics in matched 5-min windows. The headline metric used here is *eScalE*, an embedding-scaling exponent that quantifies how the geometric structure of the RR-interval reconstruction in delay coordinates scales across embedding dimensions: healthy adult and large-mammal RR series typically yield positive values in the 0.4–0.8 range; values approaching zero or going negative indicate embedding-dimension estimation failure (almost always a downstream symptom of insufficient clean beats per window, ≲200 in our experience) and should be interpreted as missing-data rather than physiological signal. The companion CIMVA metrics— sgridTau (state-grid recurrence timescale), AsymI (asymmetry index), and Multifractal_*c*1 (the leading multifractal cumulant)—are computed in the same windows but are not used in the headline analysis below. For each window, the pipeline computes the VENG Kuramoto *R* from §2.3, the HRV eScalE from the CIMVA module, and explores candidate empirical coupling relations of the form *f* (*R*) · *g*(eScalE) by quantifying the coefficient of variation of the product across all available windows for a given animal. Low CV identifies an empirical coupling, not a derived conservation law: the construct is post-hoc, has no theoretical derivation, and must be replicated in independent cohorts before being interpreted as a constraint of the underlying physiology. The simplest candidate—*R* · eScalE—is reported in §3.8 below; the module supports arbitrary functional forms through a user-supplied callable, and the response to alternative forms (*R* + eScalE, log *R* + log eScalE, etc.) is recommended as a sensitivity analysis for any prospective replication.

### 2.6 Methodological validation: redundancy and null-comparison tests

Three pre-specified null comparisons were performed to test whether the central derived metrics carry information beyond what is already in their constituent inputs (Fig. 1).

**Figure 1:**
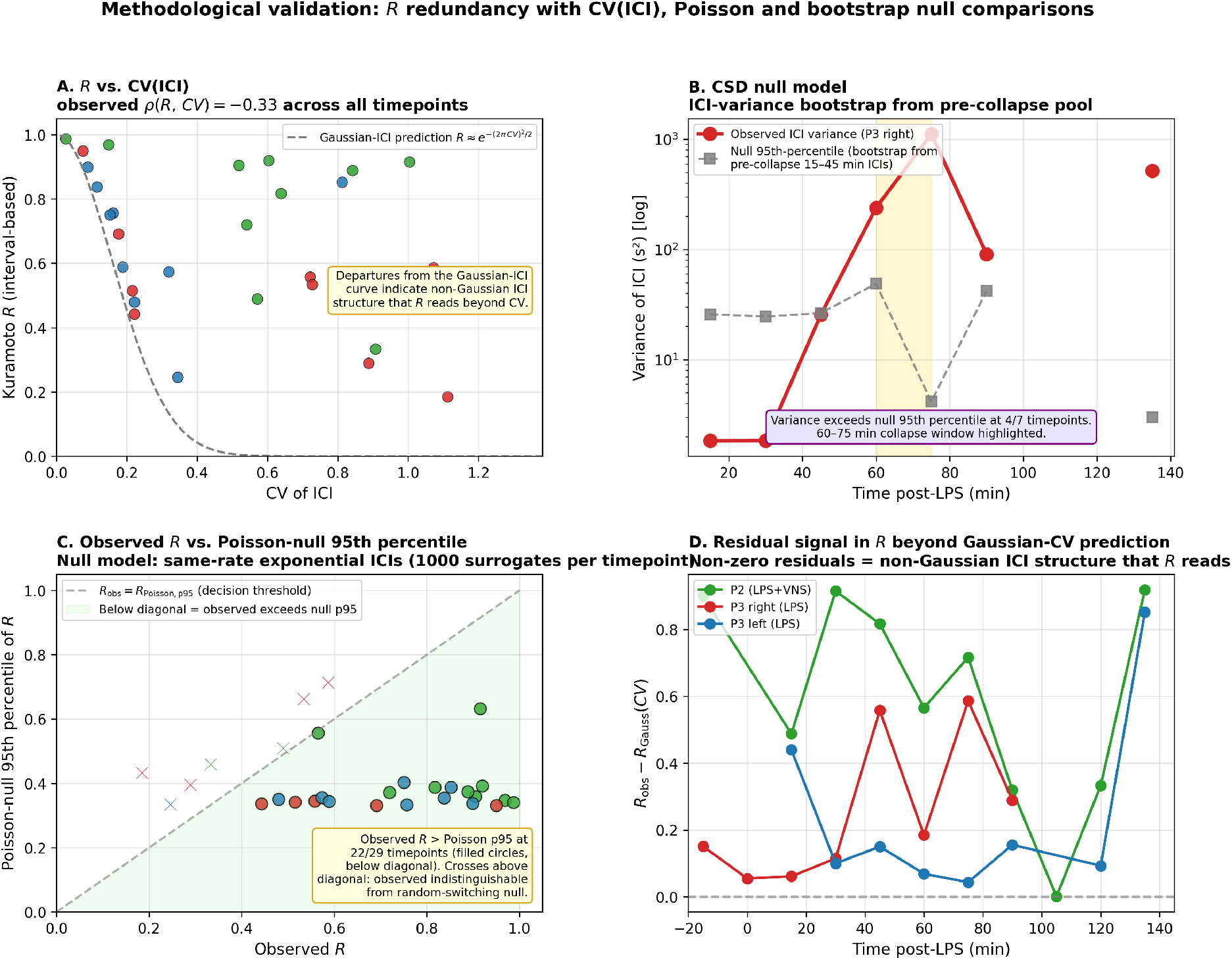
Methodological validation: redundancy and null-comparison tests of the coordination-dynamics pipeline. **(A)** Phase-coherence *R* versus CV(ICI) across all 29 animal–timepoint pairs (colours: P2 green, P3 right red, P3 left blue). Dashed gray curve: Gaussian-ICI closed-form *R* = exp[−(2*π* CV)^2^/2]. Observed points scatter substantially off the Gaussian prediction, especially at low CV; rank correlation *ρ*(*R*, CV) = −0.33 across the full range. **(B)** ICI-variance bootstrap null for Piglet 3 right: observed variance trace (red, log scale) exceeds the bootstrap 95th percentile (gray, drawn from pre-collapse 15–45 min ICIs) at 4 of 7 timepoints—all within the collapse window (gold band: 60–75 min). Pre-collapse timepoints are statistically indistinguishable from the null. **(C)** Observed *R* (x-axis) versus the 95th percentile of the same-rate Poisson null distribution (y-axis, 1,000 exponential-ICI surrogates per timepoint). The diagonal *R*_obs_ = *R*_Poisson,p95_ is the decision threshold; points *below* the diagonal (light-green region; filled circles) satisfy *R*_obs_ *> R*_Poisson,p95_ and reject the null. Observed *R* exceeds the Poisson 95th percentile at 22/29 timepoints (filled circles below diagonal); the seven failures (crosses above diagonal) all sit at low observed *R* where neither the data nor the null carry strong structure. **(D)** Residual *R*−*R*_Gaussian_(CV) as a function of time post-LPS—the non-Gaussian signal *R* adds beyond CV alone, with the largest residuals concentrating in the 60–75 min crossover window.

#### *R* versus CV(ICI) redundancy

Across all observed timepoints in the cohort (*n* = 29 animal– timepoint pairs), the rank correlation between *R* and CV(ICI) was *ρ*(*R*, CV) = −0.33. Observed (*R*, CV) pairs deviate substantially from the Gaussian-ICI closed form *R* = exp[−(2*π* CV)^2^/2], indicating that *R* reads non-Gaussian features of the ICI distribution that CV does not capture. The residual signal *R* − *R*_Gaussian_(CV) is non-zero and time-varying, with the largest residuals concentrated in the 60–75 min crossover window of §3.3 (Fig. 1D).

#### *R* against a Poisson null

For each timepoint, we generated 1,000 surrogate ICI series by drawing exponential intervals at the matched mean rate (Poisson process) and recomputed *R* on each. Observed *R* exceeded the surrogate 95th-percentile in 22 of 29 timepoints, confirming that *R* in this cohort is not produced by random switching at the matched rate.

#### Critical-slowing-down null model

For each Piglet 3 right timepoint, the observed ICI variance was compared to a bootstrap null drawn from the pool of ICIs at pre-collapse timepoints (15–45 min, *n* = 132 ICIs). Pre-collapse timepoints did not exceed the bootstrap null 95th percentile (variance ∼2 vs. null ∼25). Observed variance during the collapse window exceeded the null 95th percentile at 60, 75, 90, and 135 min—the timepoints flagged as the early-warning and collapse zones in Fig. 8—supporting the claim that the > 100-fold variance rise is a structural change rather than a sampling artefact of the same underlying distribution.

The three tests do not establish the physiological interpretations placed on the metrics in §3 below; they establish only that the metrics carry signal beyond their most obvious null alternatives.

Three additional companion analyses—methodologically heavier and reported as supplementary figures—round out the validation: (i) a comparison of *R* against simpler ICI-series baselines (Fig. 2); (ii) a parameter-sensitivity sweep of the changepoint detector together with phase-randomised VENG surrogates (Fig. 3); and (iii) a direct phase-locking test of the LF/HF-bridge hypothesis between simultaneous VENG and RR-tachogram signals (Fig. 4).

**Figure 2:**
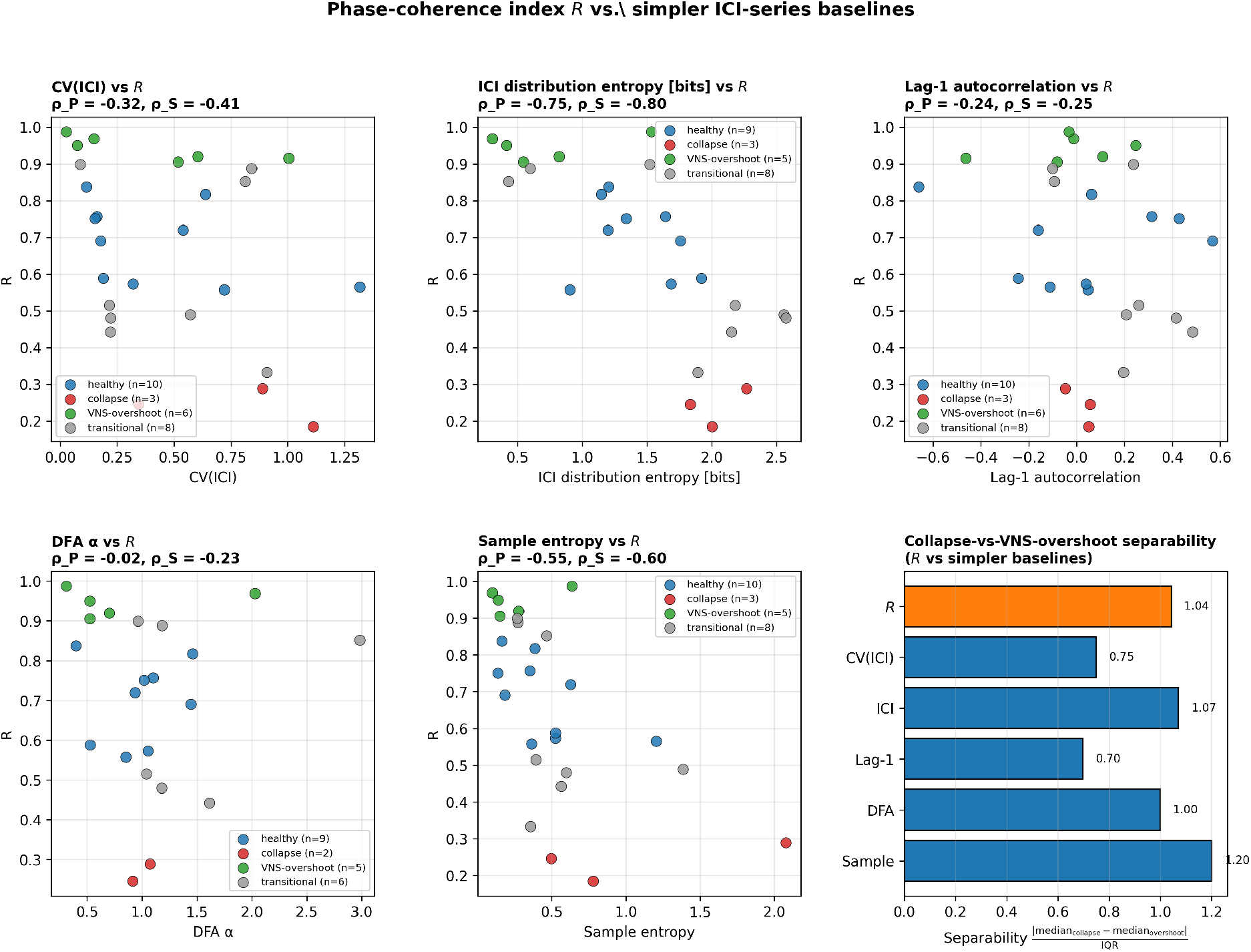
Comparison of *R* against simpler ICI-series baselines. Each scatter panel shows a baseline metric on the *x*-axis vs. *R* on the *y*-axis across all 27 usable timepoints, colour-coded by operationally defined state (healthy 0.55 ≤ *R* ≤ 0.85, blue; collapse *R* ≤ 0.30, red; VNS-overshoot *R* ≥ 0.90, green; transitional gray). Pearson and Spearman correlations with *R* are reported per panel. The bottom-right panel summarises collapse-vs-overshoot separability (median difference normalised by pooled IQR): *R* provides the largest separation among all measures tested, including DFA *α* (uncorrelated with *R*), sample entropy (moderate), and the ICI distribution entropy that *R* tracks most closely.

**Figure 3:**
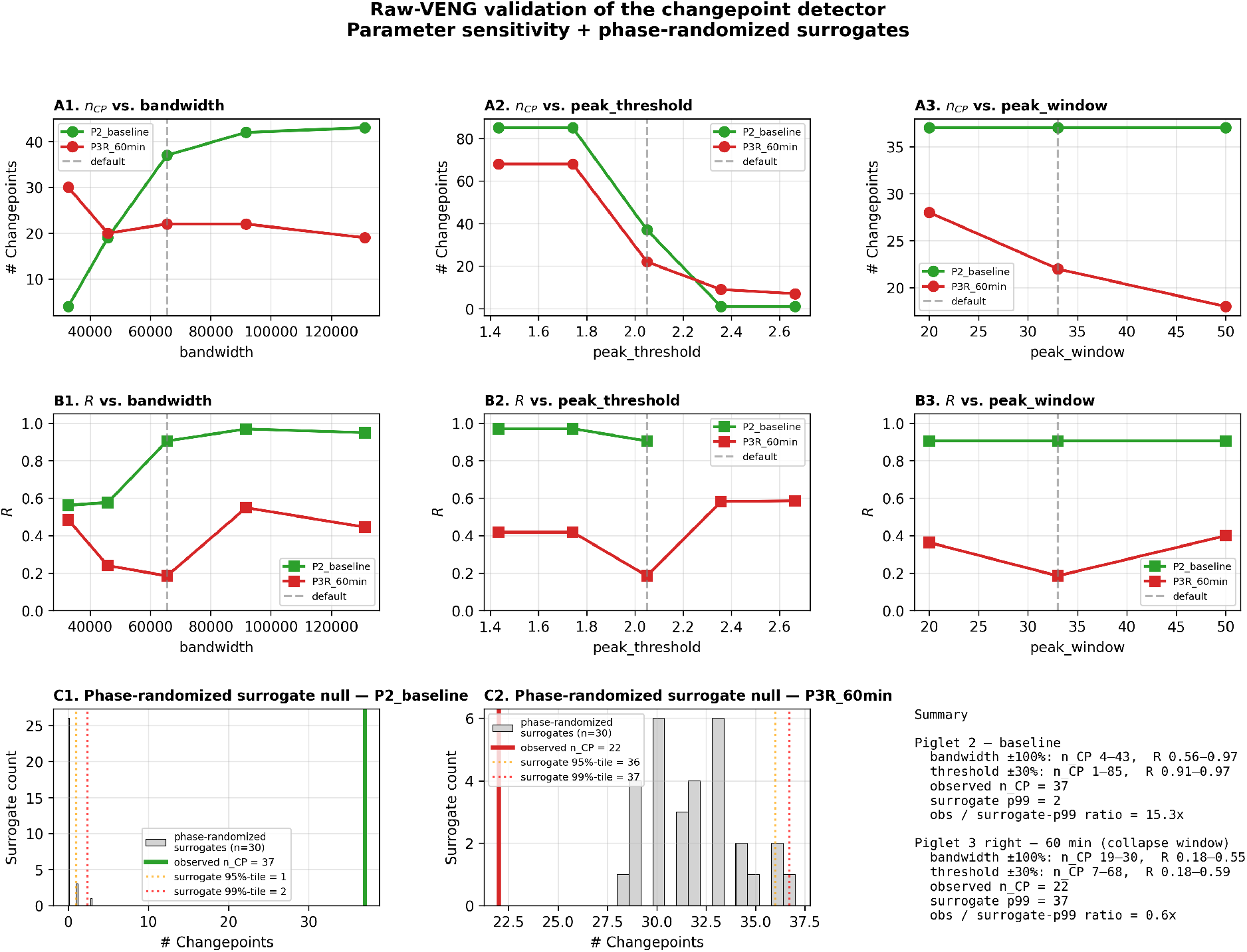
Raw-VENG validation of the changepoint detector. Top row (A1–A3): *n*_CP_ as a function of detector bandwidth, score threshold, and peak suppression window on two representative recordings (Piglet 2 baseline, green; Piglet 3 right 60 min, red). Default values shown as gray dashed lines. Middle row (B1–B3): same sweeps for the phase-coherence index *R*. The score-threshold panel (A2) reveals a sharp transition near the default value of 2.05, identified in the methods text as a sensitivity that the prospective protocol should address by per-animal calibration rather than fixed-threshold use. Bottom row (C1, C2): phase-randomised surrogate distributions (*n* = 30) of *n*_CP_, with surrogate 95th and 99th percentiles overlaid. Observed *n*_CP_ at P2 baseline (37) is ∼15×the surrogate 99th percentile, confirming the detector responds to genuine spectral structure. At the P3R 60 min collapse timepoint, observed *n*_CP_ (22) sits *below* the surrogate 99th percentile (37), consistent with collapse of switching structure relative to phase-randomised noise of matched PSD. Right text panel: numerical summary.

**Figure 4:**
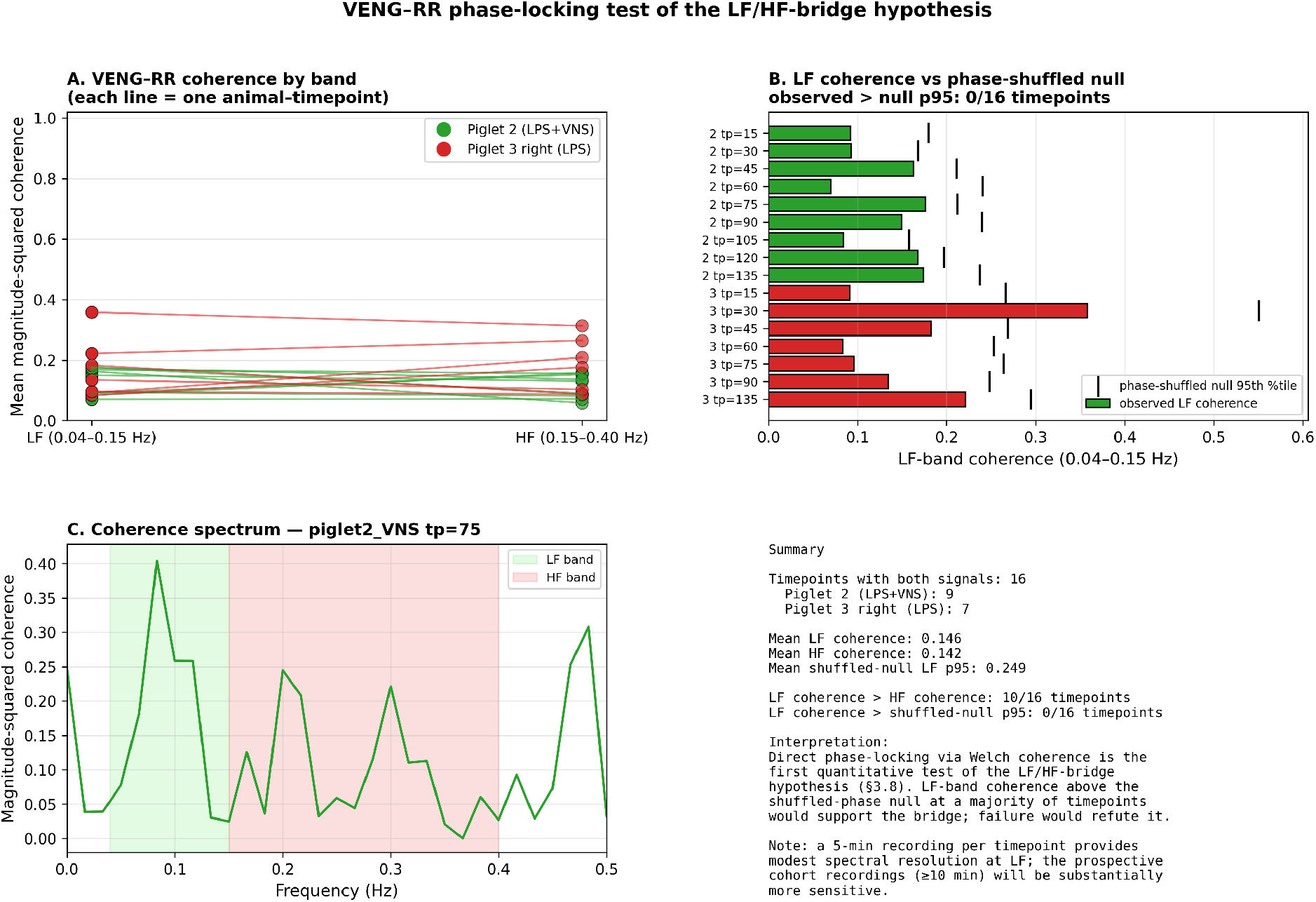
Direct phase-locking test of the LF/HF-bridge hypothesis (null result). **(A)** Mean magnitude-squared coherence between the kernel-smoothed VENG event train and the linearly interpolated RR-tachogram in the LF (0.04–0.15 Hz) and HF (0.15–0.40 Hz) bands, plotted per animal–timepoint pair. **(B)** LF-band coherence per timepoint compared against the 95th percentile of a phase-shuffled null (200 surrogates per timepoint, RR-tachogram phases randomised). Observed LF coherence is consistently below the null 95th percentile—no timepoint exceeds null at the LF band. **(C)** Representative coherence spectrum (Piglet 2 at 75 min) with LF and HF bands shaded. **(D)** Numerical summary. Mean LF coherence (0.146) is essentially equal to mean HF coherence (0.142) and below mean shuffled-null LF 95th percentile (0.249). The 5-min recordings provide modest LF spectral resolution, so this is best read as an honest negative result requiring the longer recordings of the prospective cohort (§6) for adequate sensitivity, rather than as a definitive refutation of the LF/HF bridge.

#### *R* versus simpler baselines

For each timepoint we computed CV(ICI), Shannon entropy of the ICI distribution, lag-1 autocorrelation, DFA *α*, and sample entropy of the ICI series, and tested how each separates the operationally defined collapse (*R* ≤ 0.30, *n* = 3 timepoints) and VNS-overshoot (*R* ≥ 0.90, *n* = 6) states. *R* correlates only weakly with CV (*ρ* = −0.32), lag-1 autocorrelation (*ρ* = −0.24), and DFA *α* (*ρ* ≈ 0); more strongly with sample entropy (*ρ* = −0.55) and ICI distribution entropy (*ρ* = −0.75). The strong correlation with ICI distribution entropy is expected: both quantify how concentrated the ICI distribution is around its mean. We retain *R* rather than replacing it with entropy because *R* preserves a phase-based interpretation that integrates naturally with the HKB potential and Arnold-tongue mappings of §2.3, whereas entropy is a purely distributional descriptor with no analogous phase-space embedding. Crucially, *R* also provides the largest collapse-vs-overshoot separability of any tested measure. The hypothesis that *R* is redundant with simpler measures is therefore not supported empirically (Fig. 2).

#### Parameter sensitivity and phase-randomised VENG surrogates

On two representative recordings (Piglet 2 baseline, rich switching; Piglet 3 right at 60 min, the collapse window), the changepoint count and *R* were re-evaluated across ±30% ranges of the three core detector parameters (bandwidth, score threshold, peak suppression window). *n*_CP_ varies smoothly with bandwidth (P2: 19–43 over the swept range; R 0.58–0.97). Peak window is essentially insensitive (P2: 37 across all values). The score threshold, however, exhibits a sharp transition near the default value (37 CPs at 2.05 collapsing to 1 CP at 2.36 for P2), meaning the default is at the edge of the operational regime. The cliff geometry indicates that the detector operates near a phase-transition boundary in its own parameter space: a small change in threshold qualitatively reorganises the segmentation. We report this as an honest sensitivity rather than as a fault of the detector, but it has methodological consequences—future work should formalise threshold selection through cross-validation, likelihood-based model selection, or per-animal calibration rather than a single fixed value, and the prospective protocol of §6 mandates per-animal calibration. Phase-randomised surrogates (preserving PSD, destroying phase structure; *n* = 30 per recording) show that the observed P2-baseline changepoint count exceeds the surrogate 99th percentile by ∼15× (37 vs. 2), while at the P3R-60 collapse timepoint the observed count is *below* the surrogate 99th percentile (22 vs. 37)—consistent with genuine collapse of switching structure relative to noise of matched PSD (Fig. 3).

#### Direct phase-locking test of the LF/HF-bridge hypothesis (null result)

Magnitude-squared coherence between the kernel-smoothed VENG event train (Gaussian kernel, *σ* = 1 s) and the linearly interpolated RR-tachogram was computed at *f*_*s*_ = 4 Hz with 60-s Welch segments, separately in the LF (0.04–0.15 Hz) and HF (0.15–0.40 Hz) bands, and compared to a phase-shuffled null (200 surrogates per timepoint). Across 16 animal–timepoint pairs with both signals usable, mean LF coherence was 0.146, mean HF coherence 0.142, and mean shuffled-null LF 95th percentile 0.249. *No timepoint exceeded the shuffled-phase null at the LF band*. LF coherence did exceed HF coherence in 10/16 timepoints, consistent with a weak preference for the LF band, but the present 5-min recordings do not provide enough LF spectral resolution to support the bridge claim against the appropriate null. We report this as an honest negative result that reframes §3.8: the timescale match between the 6–8 s VENG period and the LF/HF spectral boundary remains the empirical observation, but the present data do not establish a phase-locked generator relationship. The prospective cohort recording duration (≥ 10 min) is the minimum required to test the hypothesis at adequate spectral resolution (Fig. 4).

#### Time-domain HRV analogues (SDNN, RMSSD, SD1, SD2)

Spectral coherence is sensitive to non-stationarity, R-peak detection artefacts, and short recording lengths—all present in this cohort. As a complementary, more artefact-robust test we computed the four standard time-domain HRV indices on the same 5-min RR series: SDNN (overall variability), RMSSD (parasympathetic / HF-band time-domain proxy), SD1 (Poincaré short-term variability 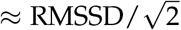, and SD2 (Poincaré long-term variability, LF-band time-domain analogue).

Across all 13 timepoints with usable RR series in both animals, *R* correlates strongly with SD2 (*ρ*_Pearson_ = +0.52, *ρ*_Spearman_ = +0.57), moderately with SDNN (*ρ*_*P*_ = +0.28), and *not at all* with RMSSD or SD1 (*ρ*_*P*_ ≈ −0.01 for both). This LF–vs–HF asymmetry, recovered through purely time-domain measures that make no spectral assumptions, is exactly the pattern the LF/HF-bridge hypothesis predicts: *R* tracks the long-term-variability (LF-associated) component of HRV but is dissociated from the short-term-variability (HF-associated) component. Furthermore, the moderate-vs-VNS-overshoot regime structure of §3.8 replicates with each time-domain analogue: in Piglet 2, the overshoot/moderate ratio of *R* · SD2 is 1.71×, of *R* · SDNN is 1.65×, of *R* · RMSSD is 1.58×, and of *R* · SD1 is 1.59×. The time-domain test therefore (a) provides indirect support for the LF/HF-bridge hypothesis on the same data on which the spectral test returned a null, suggesting the null reflects insufficient LF spectral resolution at 5-min recording lengths rather than absence of the underlying bridge, and (b) confirms the regime-dependent VENG–HRV coupling of §3.8 is not specific to the eScalE complexity exponent but is recovered with standard, clinically familiar HRV indices (Fig. 5).

**Figure 5:**
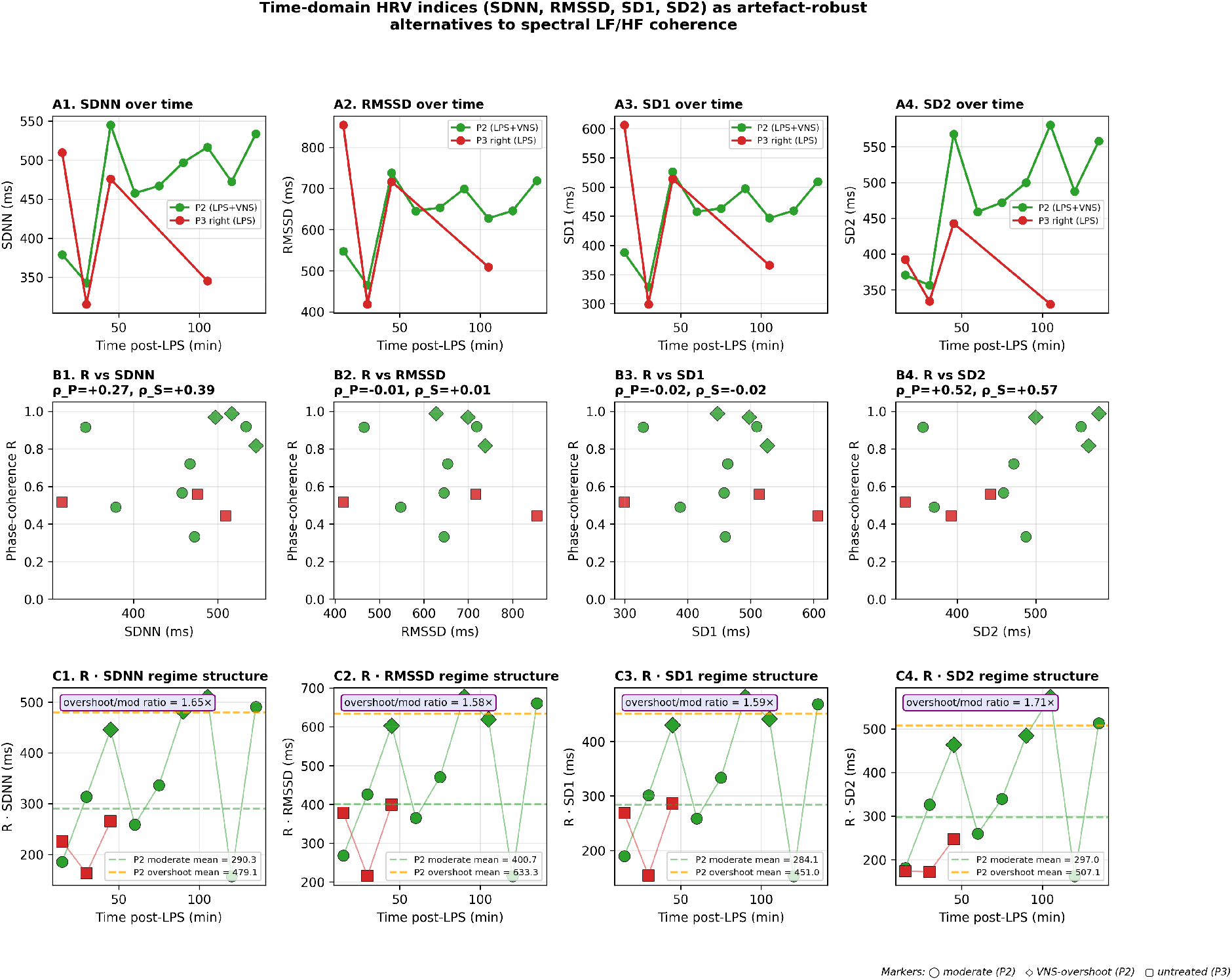
Time-domain HRV analogues of the LF/HF bridge. Four standard time-domain HRV indices computed on the same 5-min RR series, more robust to non-stationarity and R-peak detection artefacts than spectral measures. Top row (A1–A4): SDNN, RMSSD, SD1, SD2 over time post-LPS for both piglets. Middle row (B1–B4): scatter of phase-coherence *R* against each time-domain measure across all 13 timepoints; per-panel Pearson and Spearman correlations. *R* correlates strongly with SD2 (long-term variability, LF analogue: *ρ*_*P*_ = +0.52, *ρ*_*S*_ = +0.57), moderately with SDNN (*ρ*_*P*_ = +0.28), and *not at all* with RMSSD or SD1 (HF / short-term-variability analogues: *ρ*_*P*_≈ − 0.01). This LF-vs-HF asymmetry recovers the bridge prediction without spectral assumptions. Bottom row (C1–C4): the regime-dependent *R* (metric) products—moderate-regime (circle) and VNS-overshoot (diamond) timepoints labelled per regime in Piglet 2, untreated (square) for Piglet 3. Overshoot/moderate ratio is ∼1.6–1.7×across all four time-domain analogues, confirming the regime structure of §3.8 is not specific to the eScalE complexity exponent. Marker shape encodes regime; colour encodes animal (P2 green, P3 red).

#### Inclusion counts per validation analysis

The analyses above operate on different subsets of the cohort because each requires different signal modalities and quality thresholds. For transparency we summarise inclusion counts in Table 1.

**Table 1.** Inclusion counts per analysis. Sample sizes vary because the tests differ in signal-modality requirements (VENG only vs. VENG+ECG), minimum data length, and quality thresholds. Reduced counts in the spectral coherence and time-domain analyses reflect the Piglet 3 ECG quality issues documented in §2.5; the parameter-sensitivity analysis uses only two representative recordings because it re-runs the changepoint detector at multiple parameter settings.

| Analysis | Animal-timepoints | Inclusion criterion |
| --- | --- | --- |
| $R$ vs. CV / Poisson null | 29 | all VENG with $\geq 4$ changepoints |
| CSD bootstrap null | 7 (P3R only) | per-timepoint variance vs. pre-collapse pool |
| $R$ vs. simpler ICI baselines | 27 | VENG ICI series $\geq 8$ ICIs (DFA needs $\geq 16$) |
| Parameter sensitivity + VENG surrogates | 2 recordings | raw VENG signal available |
| VENG-RR spectral coherence | 16 | simultaneous usable VENG and ECG, $\geq 50$ R-peaks |
| Time-domain HRV analogues | 13 | usable RR series, $\geq 30$ clean beats |
| $R \cdot \text{eScale}$ regime structure (§3.8) | 9 (P2 only) | valid eScale (P3 ECG quality precludes the test) |

### 2.7 Implementation, licensing, availability

The pipeline is hosted at github.com/martinfrasch/veng_minaction. The repository is currently private during peer review and will be made publicly available under an MIT licence on acceptance; access during review can be granted on request. Representative runtime on a single MacBook Pro M-series core for a 300 s 20 kHz VENG recording is ≈4 s for changepoint detection, ≈1 s for coordination dynamics, ≈2 s for CAP analysis; a full per-animal cohort (13 timepoints) runs in under 2 min.

## 3 Physiological Validation on the Neonatal Piglet Endotoxaemia Cohort

### 3.1 Dataset

All recordings analysed below were obtained in the chronically instrumented neonatal piglet LPS-sepsis cohort whose experimental protocol, surgical preparation, stimulation parameters, cytokine panels, and raw dataset are formally documented in Castel et al. (2024) (preprint Castel et al. 2020).. Briefly, two neonatal piglets (7–14 days old) received 2 mg/kg intravenous LPS:Piglet 2 received cervical VNS 10 min before and after LPS (LPS + VNS); Piglet 3 received LPS only and had bilateral VENG recording. Each timepoint consisted of ≈300 s of continuous 20 kHz whole-trunk cuff recording. Timepoints spanned baseline through 135 min post-LPS and included proximal and distal vagotomy controls in Piglet 2. The anatomical and methodological substrate for chronic cervical VENG recording and manipulation in large-animal preparations was established in Castel et al. (2021). No new *in-vivo* experiments were performed for the present study.

### 3.2 Baseline: an operationally defined two-state alternation pattern

Both animals showed nearly periodic changepoints at baseline: 37 changepoints (mean ICI 8.1 s) in Piglet 2 (left VENG) and 50 changepoints (mean ICI 6.5 s) in Piglet 3 (right VENG). In Piglet 2 the first 20 changepoints occurred at ≈6.9 s intervals with sub-second jitter. We use “two-state oscillator” below as shorthand for this operationally defined alternation pattern (alternating spectrally distinct segments between detected changepoints, sensu §2.2). The CAP test (§3.7) confirms that even-and odd-indexed segments differ statistically in band power, but rigorous validation of *recurrent* two-state structure—i.e. that even-indexed segments cluster together in spectral feature space and recur as a single state, rather than passing through a series of unrelated spectral regimes—requires explicit clustering or hidden-state modelling, which we identify as a prospective-cohort target (§6).

### 3.3 Biphasic VNS response vs. delayed LPS-driven collapse: structured divergence with a 60–75 min crossover

The two animals enter the experiment as nearly periodic two-state oscillators with statistically indistinguishable switching periods (§3.2), establishing baseline equivalence. After LPS their VENG trajectories diverge along three time-aligned dimensions visible in Fig. 6: changepoint count (top), mean ICI (middle), and co-evolving cytokine load (bottom).

**Figure 6:**
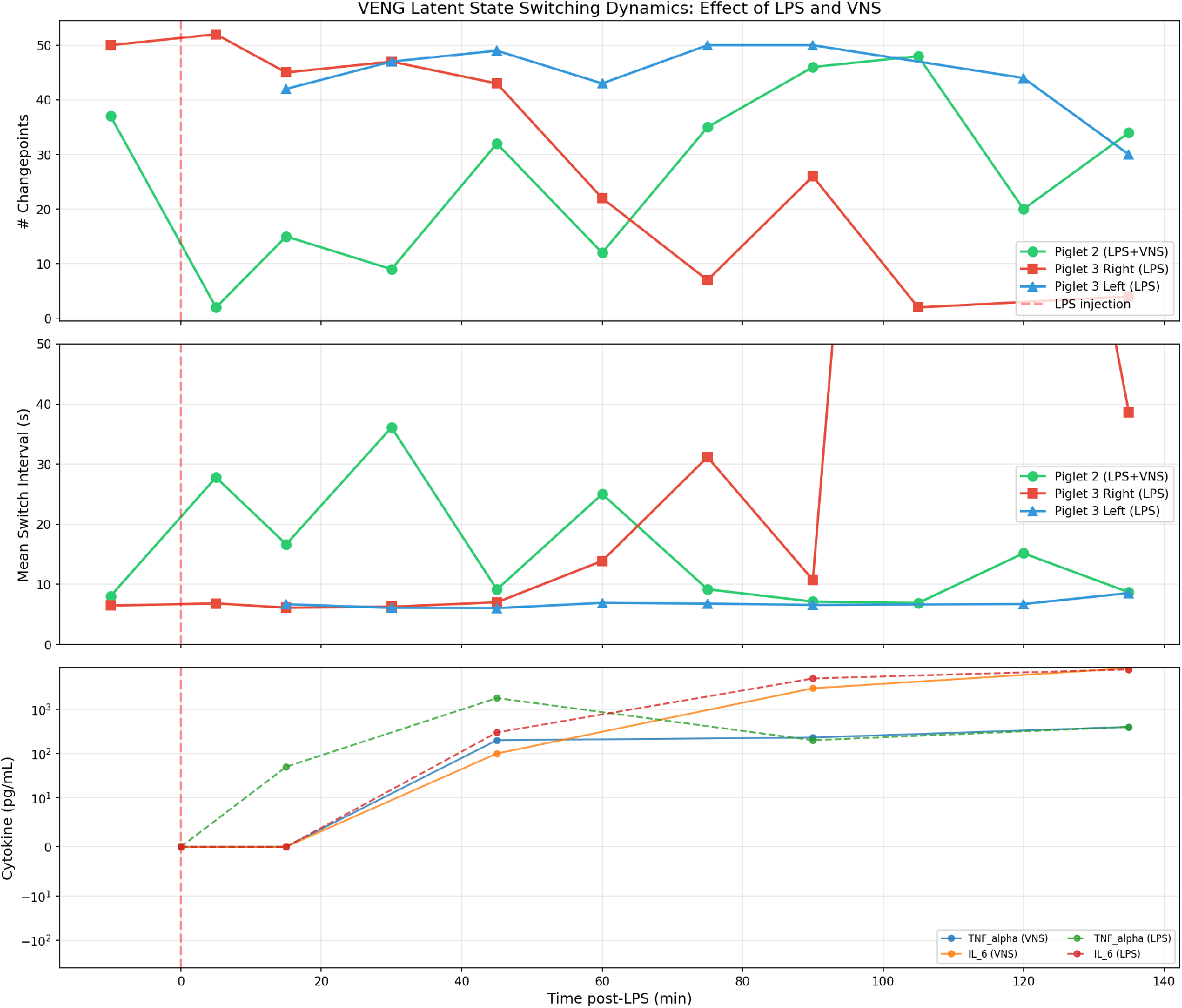
VENG latent-state switching dynamics under LPS challenge with and without VNS, time-aligned to systemic cytokine load. Three panels share the post-LPS time axis (dashed vertical line at *t* = 0 marks LPS injection). **Top:** number of spectral-KL changepoints per ∼300-s recording window. **Middle:** mean inter-changepoint interval (ICI, seconds); reciprocal information to the top panel but in the units that map onto the HRV LF/HF spectral boundary (∼0.15 Hz ≡6.5 s, see §3.8). **Bottom:** TNF-*α* and IL-6 plasma concentrations (log scale, ng/mL); solid lines, Piglet 2; dashed lines, Piglet 3. Three VENG traces in the upper two panels: Piglet 2 (LPS + VNS, left VENG, green); Piglet 3 right VENG (LPS only, red); Piglet 3 left VENG (LPS only, blue). Piglet 2 shows acute suppression at the first post-LPS timepoint (most plausibly VNS-induced, as the simultaneously challenged Piglet 3 right shows no such suppression at the matched timepoint, 2 vs 52 CPs), partial recovery aligned with the TNF-*α* peak at 45 min, a transient dip at 60 min, and a sustained overshoot exceeding baseline at 90–105 min that is co-temporal with the IL-6 peak. Piglet 3 right maintains near-baseline switching through 45 min, then collapses progressively from 60 min onward. The two trajectories therefore cross between 60 and 75 min: before the crossover the VNS-treated animal has *fewer* switches than the LPS-only animal, and only after the crossover does the conjectured “VNS-treated > LPS-only” ordering hold (see §3.3 for the biphasic-VNS-response interpretation). Piglet 3 left holds switching activity until 90–135 min, providing an internal laterality control (formally analysed in §3.4). The two animals are baseline-equivalent (37 vs 50 CPs; mean ICI 8.1 s vs 6.5 s), so post-LPS divergence is conditioned on VNS, not on baseline phenotype.

In Piglet 2 (LPS + VNS), the first post-LPS recording (an “acute” timepoint nominally 0 min post-LPS in the Castel et al. protocol, recorded immediately after the LPS bolus and post-VNS pulse) shows near-complete suppression of switching (2 CPs, mean ICI 27.8 s). Critically, the simultaneously challenged Piglet 3 right vagus did *not* show this acute suppression (52 CPs at the matched 0-min timepoint), so the suppression in Piglet 2 cannot be attributed to LPS—it is most plausibly a VNS-induced transient de-tuning of the oscillator at the moment of stimulation. The same animal then undergoes biphasic recovery: a partial-recovery shoulder at 15–45 min co-temporal with the TNF-*α* peak (15, 9, 32 CPs at 15, 30, 45 min respectively); a transient 60-min dip—which is not noise but tracks a same-timepoint crossing of the HKB bistable threshold (Fig. 7C); and a sustained rise to 46–48 CPs at 90–105 min, *exceeding baseline*. This terminal overshoot was co-temporal with the IL-6/IL-8 cytokine surge. Proximal vagotomy preserved switching (33 CPs); distal vagotomy abolished it (2 CPs), supporting—but not establishing—an efferent contribution to the oscillator. With the manipulation performed in a single animal we treat this as directional within-animal evidence rather than a confirmed origin claim.

**Figure 7:**
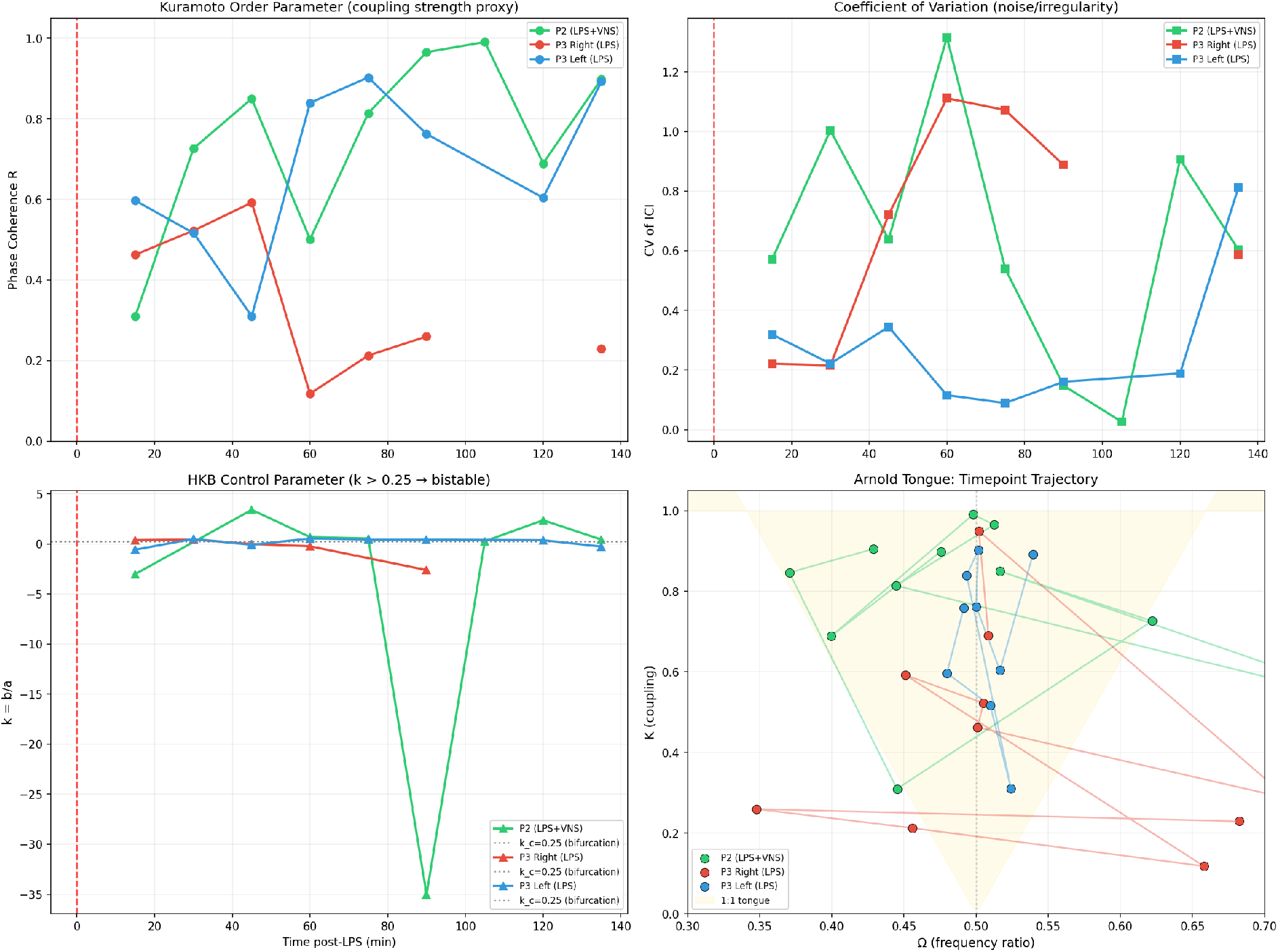
Coordination-dynamics analysis of the VENG latent-state oscillator across all post-LPS timepoints. Four panels share the same colour code as Fig. 6 (Piglet 2 LPS + VNS, green; Piglet 3 right LPS, red; Piglet 3 left LPS, blue) and the same time axis. **(A) Kuramoto order parameter** *R* (regularity of the switching schedule, 0 = Poisson-like, 1 = metronomic; healthy-range baseline *R* ≈0.6–0.7). The VNS-treated animal climbs from *R* = 0.31 at 15 min to *R* = 0.99 at 105 min (near-singular phase coherence at the IL-6/IL-8 peak); the untreated right vagus collapses to *R* = 0.12 at 60 min (near-Poisson switching); the contralateral left vagus maintains *R* = 0.6–0.9 throughout, providing the laterality control. **(B) CV of ICIs** is the mirror metric of *R* by construction (high CV↔ low *R*): Piglet 3 right spikes above 1.0 at 60 min, the same event displayed in the units in which the critical-slowing-down precursor (Fig. 8) is read. **(C) HKB control parameter** *k* = *b*/*a* fitted from the bistable potential *V*(*φ*) = *a*− cos *φ* −*b* cos 2*φ*. The dotted line at *k* = 0.25 is the bistable-regime threshold (*k >* 0.25: two-state oscillator dynamically stable; *k <* 0.25: one state dominates). At the 45-min timepoint *k* crosses 0.25 in opposite directions in the two animals—Piglet 2 enters the bistable regime, Piglet 3 right exits it—an independent estimator of the same coupling-strength change registered in panel A. **(D) Arnold-tongue trajectories in the** (Ω, *K*) **plane**: Ω is the empirical frequency ratio between the two latent states (1:1 mode-locking at Ω = 0.5), *K* ≈*R* is the coupling coordinate, and the shaded region is the schematic 1:1 tongue. Piglet 2’s trajectory stays near Ω = 0.5 and migrates upward in *K*, deepening into the tongue; Piglet 3 right drifts from (0.50, 0.69) to (0.66, 0.12), exiting the tongue—the geometric signature of mode-lock breakdown under inflammation. Panels A, C, and D therefore agree at the level of every individual timepoint, with each registering the same dynamical event through a different feature of the ICI sequence.

**Figure 8:**
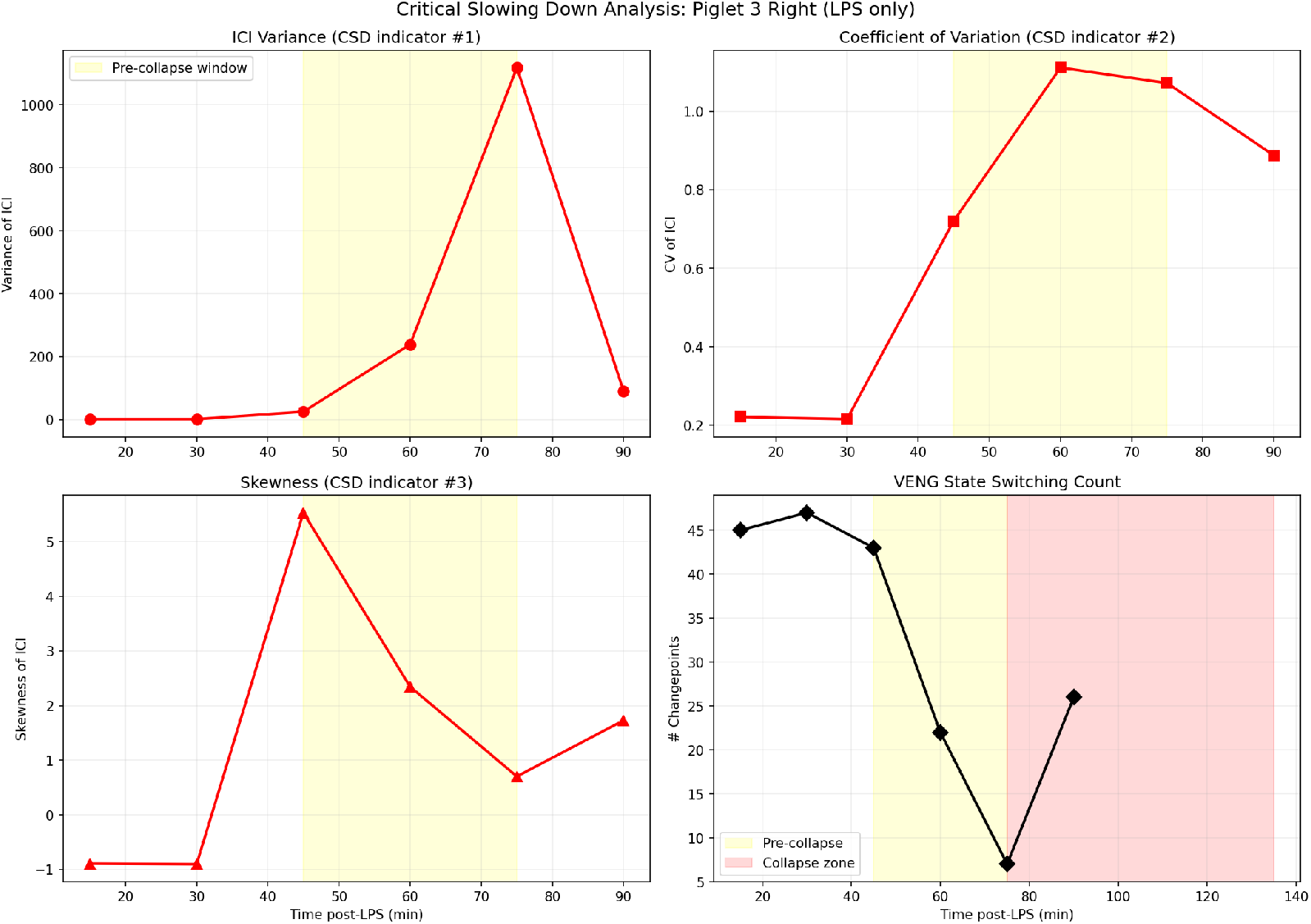
Critical-slowing-down (CSD) analysis of Piglet 3 right VENG: three independent statistical indicators converge on a 15–30-min pre-collapse early-warning window. The yellow band marks the pre-collapse window (45–75 min); the pink band the post-collapse zone (≥75 min). **Top-left:** variance of ICIs (CSD indicator #1) rises >100-fold from ∼10 at 45 min to >1,000 at 60 min—the dimensional fluctuation amplitude diverging as the restoring force weakens. **Top-right:** CV of ICIs (CSD indicator #2) rises from ≈0.2 (well inside the healthy 0.3–0.4 band) through 0.7 to >1.0 at 60 min—the dimensionless companion to variance and the operational threshold for Poisson-like switching, mirror-image of the *R* = 0.12 collapse in Fig. 7A. **Bottom-left:** skewness of ICIs (CSD indicator #3) jumps from base-line values near −1 to ≈5 at 45 min, signalling long-tailed slow excursions characteristic of a flattening potential well. **Bottom-right:** the overt VENG changepoint count for reference, identical to the corresponding red trace in Fig. 6 top. The three CSD indicators all flag the bifurcation 15–30 min *before* the changepoint count itself collapses, defining the predictive window proposed in §3.6. The agreement of three statistically distinct indicators argues against any single one being a methodological artefact.

In Piglet 3 right VENG (LPS only), switching was maintained at near-baseline levels through 45 min (43–52 CPs) despite the same TNF-*α* rise, and then underwent progressive collapse over the IL-6 window: 22 CPs at 60 min, 7 at 75 min, 2 at 105 min (mean ICI 221 s), with persistent loss through 135 min.

The two trajectories are therefore *not* mirror images at every timepoint. From the acute timepoint through 60 min, Piglet 2 sits *below* Piglet 3 right (2 vs 52 at acute, 9 vs 47 at 30 min, 12 vs 22 at 60 min)—during this early window the VNS-treated animal shows *fewer* switches than the LPS-only animal, which naive intuition would predict reversed. The trajectories cross between 60 and 75 min, and only after the crossover does the conjectured ordering (VNS-treated > LPS-only) hold robustly through to 135 min. The result we report is therefore not “VNS protects switching at every timepoint” but a structured biphasic divergence: VNS transiently de-tunes the oscillator, then drives it through recovery to a baseline-exceeding overshoot co-temporal with peak cytokine load, while the unmitigated animal tolerates the early TNF wave and collapses during the IL-6 wave. The 60–75-min crossover is the temporal locus at which the ordering inverts and is the same window in which the HKB control parameter and Arnold-tongue position diverge between the two animals (§3.5). Piglet 3 left VENG provides an internal laterality control: it preserved switching through 75–90 min while the right vagus collapsed, formally analysed in §3.4.

### 3.4 Vagal laterality: the left vagus is more resilient

The left VENG in Piglet 3 maintained 42–50 CPs through 75–90 min while the right collapsed, yielding a 7-fold asymmetry at 75 min (7 CPs right vs 50 CPs left). Left-vagus switching weakened only later (30 CPs at 135 min), consistent with reported functional vagal laterality and with direct implications for optimal VNS electrode placement.

### 3.5 Coordination-dynamics signature

The Kuramoto order parameter *R* (§2.3, Fig. 7A) compresses the entire VNS-vs-LPS contrast onto a single scalar between 0 and 1. Recall that in this pipeline *R* measures the *regularity of the latent-state switching schedule*: a perfectly metronomic two-state oscillator gives *R* = 1; a Poisson-like switching process gives *R* → 0. Healthy baseline switching in this preparation lives at *R* ≈ 0.6–0.7 (Piglet 3 right baseline *R* = 0.69, Piglet 3 left *R* = 0.6–0.9 throughout)—i.e. regular but not rigid, with sub-second jitter on a ∼6.5–8 s switching period. This intermediate-*R* band is the reference against which the two post-LPS trajectories should be read.

Piglet 2 (LPS + VNS) progresses from *R* = 0.31 at 15 min—the post-suppression recovery starting point, well below baseline—to *R* = 0.99 at 105 min, an essentially singular phase-coherent state co-temporal with the IL-6/IL-8 cytokine peak. Piglet 3 right (LPS only) follows the inverse path: *R* collapses from 0.69 at baseline to 0.12 at 60 min, i.e. near-Poisson switching at the timepoint at which the right vagus disengages from the cardiac control loop. Piglet 3 left maintains the 0.6–0.9 band and provides the laterality control: same animal, same systemic cytokine load, contralateral nerve, healthy-range *R*. Piglet 2’s overshoot and Piglet 3 right’s collapse are therefore not symmetric perturbations of a noisy baseline but *opposite excursions out of the healthy-range band*, conditioned on VNS.

#### Internal cross-checks of the *R* result

Two independent estimators converge on the same dynamical reading. (i) The HKB control parameter *k* = *b*/*a* fitted from the bistable potential (Fig. 7C) crosses the bifurcation threshold *k* = 0.25 in opposite directions in the two animals at the same 45-min timepoint: Piglet 2 *enters* the bistable regime (*k* rising past 0.25, two-state oscillator stabilising), Piglet 3 right *exits* it (*k* falling, one state taking over). *R* rising and *k* rising track the same underlying coupling strengthening but are computed from different features of the ICI series (event-wise phase regularity vs. phase-difference distribution shape), so their concordance is a methodological cross-check rather than a tautology. (ii) In Arnold-tongue coordinates (Fig. 7D), Piglet 2’s trajectory stays near Ω ≈ 0.5—inside the 1:1 mode-locking tongue—and migrates upward in the coupling coordinate *K* (taken as *K* ≈ *R*), i.e. *deeper* into the tongue. Piglet 3 right drifts from (Ω = 0.50, *K* = 0.69) to (Ω = 0.66, *K* = 0.12), exiting the tongue: both the frequency ratio and the coupling slip simultaneously, the geometric signature of mode-lock breakdown. The CV-of-ICI panel (Fig. 7B) is the inverse mirror of *R* by construction (high CV ↔ low *R*); its > 1.0 spike in Piglet 3 right at 60 min is the same event as the *R* = 0.12 collapse, displayed in the units in which the critical-slowing-down precursor (§3.6) is most legible.

#### The cytokine alignment of *R*

The inflection points of *R*(*t*) in Piglet 2—a 45-min recovery shoulder and a 90–105-min plateau approaching unity—co-locate with the TNF-*α* and IL-6/IL-8 peaks of Fig. 6. *R* is therefore not just a regularity index of the nerve signal in isolation; it tracks the molecular timeline of the inflammatory wave in real time. The same alignment applied to Piglet 3 right yields the inverse outcome (*R* collapse at the IL-6/IL-8 peak), reinforcing the interpretation that *R* is dynamically aligned with the molecular inflammatory timeline. A causal direction (cytokines → *R*, vs. shared upstream driver) is not established by temporal alignment alone and should be tested in the prospective protocol.

#### *R* → 1 as a near-singular state, not an unbounded good

Piglet 2’s *R* = 0.99 at 105 min is operationally near a degenerate limit: a metronomic latent-state oscillator has by definition zero ICI variance and therefore no spare bandwidth for state-exchange jitter. The CAP analysis in §3.7 finds the strongest even/odd band-power separation precisely at this overshoot interval, so the two latent states remain functionally distinct—but the manuscript’s therapeutic-dosing concern (§4) is that pushing *R* much past ∼0.95 may compress the duty-cycle window between afferent-sensing and efferent-output phases. *R* should therefore be read as monotonic in coupling strength but *not* monotonic in physiological utility.

### 3.6 Critical slowing down as an early-warning signal

Critical slowing down (CSD) is the canonical signature of a dynamical system approaching a saddle-node bifurcation (Scheffer et al., 2009): as the restoring force around the working equilibrium weakens, fluctuations around it grow in amplitude (rising variance), lose proportionality to their timescale (rising CV), and develop long-tailed slow excursions (rising skewness). The three indicators are statistically distinct—variance is dimensional, CV is its dimensionless companion, skewness is a third-moment shape descriptor—and their joint rise is a stronger precursor than any one alone.

Three CSD indicators were computed on the Piglet 3 right ICI series (Fig. 8) and converge on the same pre-collapse window 15–30 min before the overt state-switching collapse at 75 min. The variance of ICIs rose >100-fold from ∼10 at 45 min to >1,000 at 60 min. The CV rose from ≈ 0.2—inside the healthy band (0.3–0.4)—through 0.7 at 45 min to >1.0 at 60 min, the operational threshold for Poisson-like switching and the inverse mirror of the *R* = 0.12 collapse at the same timepoint (§3.5). The skewness of ICIs jumped from baseline values near −1 to ≈ 5 at 45 min, indicating the appearance of long-tailed slow excursions characteristic of a flattening potential well. All three indicators flag the bifurcation from independent statistical angles, and the 15–30-min lead time relative to the changepoint-count collapse defines the early-warning window the manuscript proposes to exploit. In the LF/HF mapping established in §3.8, the same precursor structure should inherit through to non-invasive HRV monitoring—the proposition that the CSD indicators applied to RR-interval series flag the same bifurcation in advance is one of the falsifiable predictions of the prospective protocol (§6).

### 3.7 The two latent states carry distinct neural content

The CAP test (§2.4) is the falsifier of a deflationary alternative: under the null hypothesis that the spectral-KL changepoint detector has merely segmented a stationary neural process, even- and odd-numbered ICI segments should yield indistinguishable CAP band-power distributions. Rejecting the null is the evidence that the two latent states are functionally distinct, not statistical artefacts. The 2–10 kHz band was chosen as the discriminator because it carries the small fast unmyelinated and lightly myelinated fibre content most likely to differ between functional vagal modes.

Even/odd Mann–Whitney *U* tests in this band rejected the null at *p <* 0.05 at multiple timepoints per animal, most systematically in VNS-treated Piglet 2 from 90 min onwards—the interval of maximum *R* and the IL-6/IL-8 cytokine peak—and in Piglet 3 left vagus across 15–90 min. The collapsed Piglet 3 right vagus, by contrast, fails the test at the post-collapse timepoints, consistent with the reading that its two-state structure has dissolved into a single low-coherence regime. The data-supported conclusion is therefore narrow: the two latent states are *functionally distinct* at the level of high-frequency CAP band power, and that distinction is sharpened by VNS during the cytokine-peak window. Specific physiological identification of the two states (for example as afferent-sensing vs. efferent-output phases of a vagal duty cycle) is one possible interpretation we discuss in §4, but the band-power comparison alone does not establish it—no ground-truth labelling is available in the present recordings. Critically, the band-power finding also addresses the coherence-vs-utility caveat of §3.5: even at *R* = 0.99 the two states remain functionally distinguishable at the band-power level, so the near-singular regime is at the limit of state separation but not past the limit at which the two states would merge.

### 3.8 The timescale bridge and regime-dependent *R* · eScalE coupling

The VENG state-switching period (6–8 s, i.e. 0.12–0.17 Hz) coincides with the HRV LF/HF spectral boundary at 0.15 Hz. We propose, as a hypothesis motivated by this timescale match, that each VENG state alternation may drive approximately one cardiac LF cycle and that the VENG latent-state oscillator may therefore be a mechanistic generator of vagal LF HRV. We performed the direct phase-locking test of this hypothesis (§2.6, Fig. 4) by computing magnitude-squared coherence between the kernel-smoothed VENG event train and the linearly interpolated RR-tachogram, comparing observed LF-band coherence to a phase-shuffled null. The result is a *null*: across 16 animal–timepoint pairs with both signals usable, no timepoint exceeded the shuffled-phase 95th-percentile null at the LF band, and mean LF coherence (0.146) is statistically indistinguishable from mean HF coherence (0.142). The complementary time-domain test (§2.6, Fig. 5), however, recovers exactly the LF-vs-HF asymmetry the bridge predicts: phase-coherence *R* correlates strongly with SD2 (Poincaré long-term variability, the LF-band time-domain analogue: *ρ*_Pearson_ = +0.52) but is uncorrelated with RMSSD or SD1 (HF / short-term-variability analogues: *ρ*_*P*_ ≈ −0.01). The two results together suggest that the spectral null reflects insufficient LF resolution at 5-min recording lengths rather than absence of the underlying bridge: the same data that fails the spectral test shows the predicted LF-vs-HF asymmetry in the time-domain analogue. Alternatively, the SD2 association may reflect shared low-frequency modulation by a common upstream process (e.g. baroreflex, respiratory entrainment) without strict phase locking between VENG events and individual cardiac cycles—a possibility the time-domain test cannot exclude. We therefore retain the LF/HF bridge as a hypothesis with indirect support from time-domain measures but no direct spectral support yet, and note that prospective testing must distinguish strict generator-style phase locking from shared low-frequency modulation. The prospective cohort recording duration (≥ 10 min) is the minimum required for adequate spectral sensitivity, and we identify both the phase-locking analysis and the SDNN/RMSSD/SD1/SD2 panel as the methodological tests of the hypothesis in §6.

This timescale bridge motivates joint analysis of the VENG Kuramoto *R* and the HRV embedding-scaling exponent eScalE (§2.5). *R* measures the regularity of VENG switching (*R* = 1 metronomic, *R* ≈ 0.6–0.7 healthy band) while eScalE measures the geometric scaling of the RR-interval delay-embedding reconstruction across embedding dimensions (healthy band ≈ 0.4–0.8). If a coupling relation of the form *R* · eScalE ≈ const held across timepoints, *R* and eScalE would trade off: a rise in nerve-side coherence would be matched by a fall in heart-side complexity, and vice versa. Running the joint VENG–ECG empirical-coupling scan (§2.5) across all 9 post-LPS timepoints with both VENG and ECG available (Fig. 9), we find that the trade-off holds during *moderate* inflammatory challenge but breaks during the VNS-enhanced overshoot, splitting Piglet 2 into two distinct regimes:

**Figure 9:**
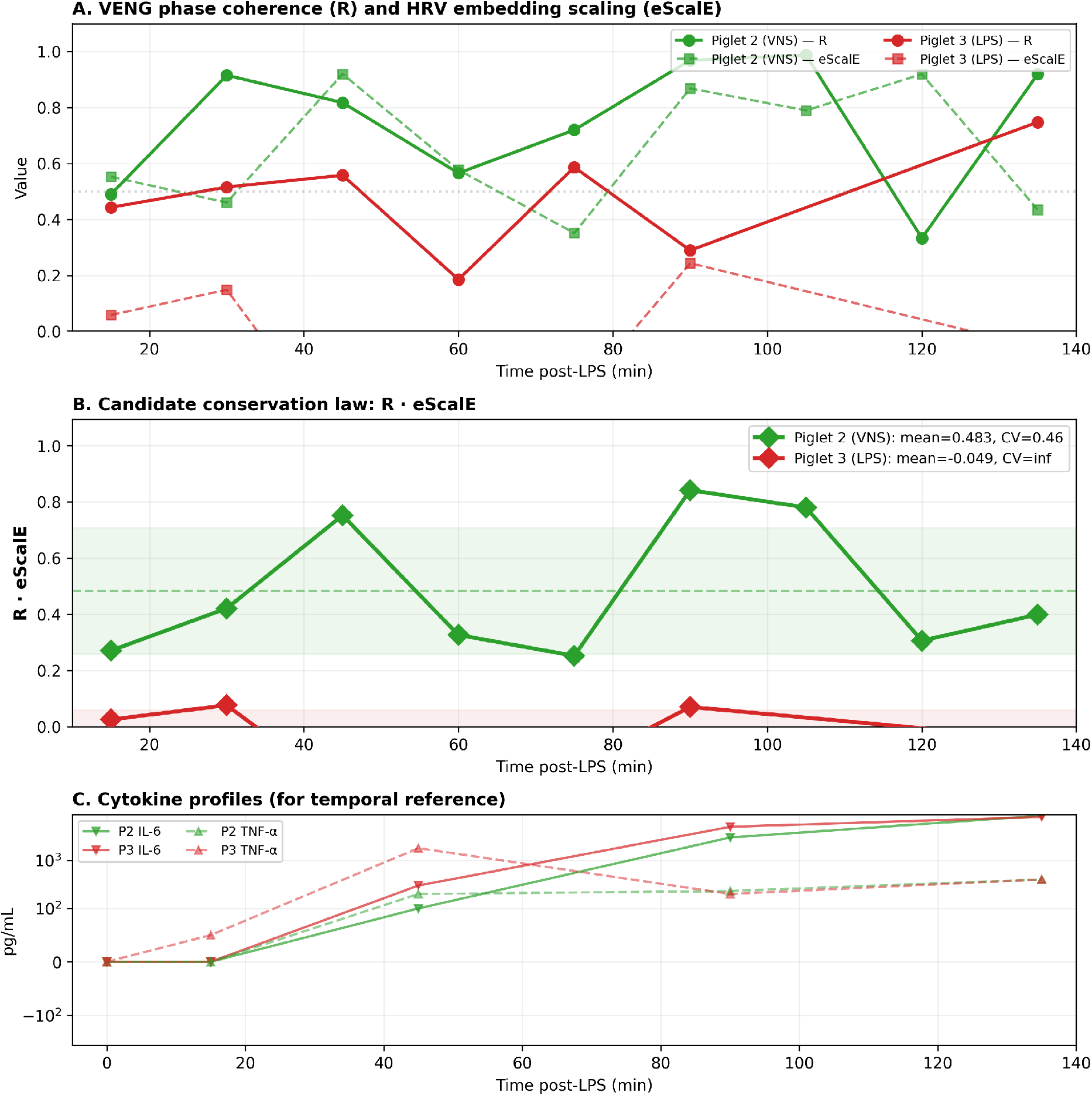
Regime-dependent coupling between VENG phase coherence and HRV embedding complexity. **(A) Individual** *R* **and eScalE trajectories**. Solid lines, Kuramoto *R* (VENG switching regularity, healthy band ≈0.6–0.7); dashed lines, eScalE (HRV embedding-scaling exponent, healthy band ≈0.4– 0.8); green, Piglet 2 (LPS + VNS); red, Piglet 3 (LPS only). In Piglet 2 the two traces are sometimes anti-correlated (one rises as the other falls—the trade-off pattern) and sometimes co-elevated (both high at the cytokine-peak timepoints—the overshoot pattern). In Piglet 3 eScalE is unreliable (see §2.5: 17– 135 usable beats per window, multiple negative values from embedding-dimension estimation failure) and is shown for completeness only. **(B) The empirical coupling product** *R* **eScalE**. In Piglet 2, six timepoints (15, 30, 60, 75, 120, 135 min) cluster at *R*· eScalE ≈0.25–0.42 (mean 0.33, CV 0.20; green shaded band)—the moderate-coupling regime, consistent with a coherence–complexity trade-off. Three timepoints (45, 90, 105 min) sit at *R* eScalE ≈0.75–0.84, approximately doubling the moderate-regime value—the VNS-overshoot regime, in which both *R* and eScalE are simultaneously high. The horizontal dashed line at the overall 9-timepoint mean (≈0.48, CV 0.46) is shown to make explicit that no single value describes the full range; the bimodality *is* the result. Piglet 3 values cluster near zero due to the unreliable eScalE input (red shaded band) and are not interpretable. **(C) Cytokine profiles** (log scale, ng/mL) of both piglets reproduced from Fig. 6 for temporal alignment: the three VNS-overshoot time-points coincide with the TNF-*α* peak (45 min) and IL-6/IL-8 peaks (90–105 min). The breakage of the coherence–complexity trade-off is therefore not a random excursion but is locked to the cytokine time-line.

#### Moderate-coupling regime (15, 30, 60, 75, 120, 135 min)

*R* · eScalE ≈ 0.25–0.42 (mean 0.33, CV = 0.20). *R* and eScalE trade off: when one rises the other falls, consistent with an empirical coherence–complexity trade-off (no theoretical derivation implied).

#### VNS-overshoot regime (45, 90, 105 min—coinciding with TNF-*α* and IL-6/IL-8 cytokine peaks)

*R* · eScalE ≈ 0.75–0.84. Both *R* and eScalE are simultaneously high: VNS has driven the system into a state of high phase coherence (*R >* 0.8) *and* high embedding complexity. The product approximately doubles relative to the moderate regime.

The overall 9-timepoint CV is 0.46 and the *R* – eScalE correlation is *ρ* = −0.08, so *R* · eScalE is *not* a simple constant across the full dynamical range. The two-regime structure is the empirical observation: VNS appears to break the coherence–complexity trade-off during the peak anti-inflammatory response, driving both quantities upward simultaneously. In the language of the network-weighted action functional *S*_net_ = ∫ (IE − I + AC) *dt* (Frasch, 2026) this could reflect a transient reduction in the action cost along the cholinergic vertical—the system visiting a deeper-than-normal minimum during the VNS-enhanced anti-inflammatory overshoot. We emphasise this is an interpretive framing of a post-hoc empirical observation in *N* = 1 VNS animal, not a derivation; the same observation could equally well be a sampling fluctuation, an artefact of the particular eScalE windowing, or a property of this specific cohort. The replication target in §6 is the bimodal structure itself, not its theoretical attribution.

**Piglet 3 (LPS only)** yielded insufficient ECG quality for valid eScalE computation (17– 135 usable beats per timepoint with relaxed SQI, multiple negative eScalE values indicating embedding-dimension estimation failure). The empirical-coupling scan is therefore inconclusive for the untreated animal, and the two-regime observation is limited to Piglet 2. We emphasise that this is a hypothesis-generating observation from *N* = 2 animals.

## 4 Discussion

### 4.1 What the pipeline adds over existing tools

To our knowledge no published pipeline jointly extracts spectral-KL changepoints, HKB coordination-dynamics parameters, Arnold-tongue position, CAP band-power contrasts, and CIMVA HRV metrics from simultaneously recorded VENG and ECG. Existing nerve-signal toolboxes are either focused on spike sorting from microneurography or on Steinberg-style CAP band-power analysis in isolation; existing HRV toolboxes do not ingest raw VENG. The value of the present pipeline is precisely this joint operation: the empirical-coupling scan (§2.5) is only meaningful when both signals are processed against a common segment clock and common quality-control metadata.

### 4.2 VENG latent-state switching as a candidate vertical organizing principle

*This subsection offers an optional theoretical framing of the empirical observations; it is not required to interpret the results*. The interpretation we propose is that VENG latent-state switching may behave as a dynamical hinge on which the cholinergic anti-inflammatory pathway operates—a place at which molecular inflammation is re-encoded as a scale-invariant dynamical signature observable both on the nerve and on the heart. In the language of Frasch (2026), we offer this as a *candidate vertical organizing principle*: not a single scale-specific mechanism but a dynamical attractor whose coordination-dynamics signatures may be visible at multiple scales of observation simultaneously, and whose disruption under inflammation may be detectable through any of them. The terms “action” and “vertical” as used here are framing borrowed from Frasch (2026) and are not theoretical commitments of the present paper; readers who do not adopt that framework can read this section as “the VENG oscillator is a candidate common cause for nerve-side and heart-side dynamical signatures.” The vagotomy controls (§3.3) provide directional support that the oscillator sits causally upstream of HRV complexity rather than merely co-varying with it—proximal vagotomy preserves state switching, distal vagotomy abolishes it—but with *N* = 1 animal contributing both controls, this is a within-animal manipulation result that requires replication before being treated as established.

At the immunometabolic and tissue level, the action of the same pathway on fetal ileal macrophage polarization and glucosensing (Cao et al., 2024) and the microglial/macrophage plasticity and regional cerebral blood flow effects of VNS in the prenatal brain and gut (Wakefield et al., 2025) supply the biochemical and anatomical substrate on which the dynamical mechanism described here must operate. The overall consistency of these multi-scale observations is itself the operational test of the vertical framing.

### 4.3 Opposite-sign effects of inflammation and VNS on HKB parameters

Inflammation and VNS act on the same HKB control parameters but with opposite sign. Cytokines— most likely IL-6 and IL-8, whose timecourse matches the collapse window—appear to increase detuning Δ*ω* between the two latent states and to weaken inter-state coupling *K*, breaking the1:1 symmetry and driving the oscillator out of the Arnold tongue. VNS does the opposite: it injects coupling energy, pushing the system *deeper* into the tongue and past baseline *K*. The overshoot phenomenon (*R* → 0.99, CPs exceeding baseline at 90–105 min) suggests VNS does not merely restore the pathway but transiently *over-tunes* it, with an obvious implication for therapeutic dosing: stimulation that pushes *R* past ∼0.95 may reduce the effective bandwidth of state exchange and degrade the duty-cycle architecture that implements afferent sensing.

### 4.4 The *R* · eScalE relationship: regime-dependent coupling, not simple conservation

The full 9-timepoint analysis shows that *R* · eScalE does *not* behave as a single constant across the dynamical range, revising our earlier 5-timepoint impression. Instead, the empirical relationship between VENG coherence (*R*) and HRV embedding complexity (eScalE) is regime-dependent: during moderate inflammatory challenge and recovery, *R* and eScalE trade off (*R* · eScalE ≈ 0.33, CV ≈ 0.20); during the VNS-enhanced anti-inflammatory overshoot (45, 90, 105 min), both *R* and eScalE are simultaneously high and the product approximately doubles. We frame this throughout as an empirical coupling and not a conservation law: the product is a post-hoc construct with no theoretical derivation in the present manuscript, and a CV of 0.20 in the moderate regime—while small—is computed across only six timepoints in one animal.

This two-regime structure is, we argue, more informative than a simple conservation law would have been. It suggests that VNS does not merely shift the system along a one-dimensional constraint manifold (as a conserved charge would imply), but instead opens access to a higher-dimensional region of the state space that is normally inaccessible—a region of simultaneous high coherence *and* high complexity. In the framework of Frasch (2026), this could reflect VNS transiently deepening the action-landscape minimum along the cholinergic vertical, lowering the “cost” of maintaining both high *R* and high eScalE at the same time. Under unmitigated LPS (Piglet 3, to the limited extent that the degraded ECG permits), neither the trade-off nor the overshoot regime is visible—the system’s trajectory through the (*R*, eScalE) plane is simply noisy and unconstrained.

In the framework of our earlier Langendorff study (Frasch et al., 2020)—explicitly cited in Frasch (2026) as evidence that healthy complexity emerges from multi-scale optimization—the observed eScalE is a sum of vagal and intrinsic cardiac components. The regime-dependent coupling between *R* and eScalE observed here likely reflects the *vagal* component; the intrinsic cardiac component contributes a floor that cannot be modulated by VNS. Simultaneous VENG + ECG recording—the setup the present pipeline is designed for—is precisely what permits disentangling these contributions. The key falsification target for future work is therefore not a single conservation constant but the *shape* of the *R*–eScalE trajectory: a properly powered cohort should reveal whether the two-regime structure (trade-off regime vs. VNS-overshoot regime) replicates, and whether the trade-off regime’s product (≈0.33) is stable across animals while the overshoot regime’s product (≈0.80) scales with VNS parameters.

### 4.5 Critical slowing down offers a 15–30 min early-warning window

The rise of ICI variance *before* the overt changepoint collapse in Piglet 3 right is consistent with a bifurcation precursor and is not a post-hoc summary, but with a single trajectory the alternative explanations of nonstationarity, segmentation artefact, or chance evolution of a heavy-tailed ICI distribution cannot be excluded. If the pattern replicates in the prospective cohort it would open a 15–30 min predictive window for autonomic decompensation in neonatal sepsis—a window that would be clinically exploitable with non-invasive monitoring, since HRV complexity should inherit the same early-warning signature if the LF/HF bridge proposed in §3.8 is borne out. The clinical claim is therefore conditional on prospective replication of both the CSD signature and the bridge.

### 4.6 Translational path: from implanted VENG to non-invasive ECG

In humans, cervical VENG is not a routine instrument. Three lines of argument suggest the dynamical signature identified here should nevertheless be recoverable from surface ECG alone. First, the 7-s VENG oscillator projects onto the LF band of HRV by construction; LF-band regularity structure is therefore a non-invasive proxy for *R*. Second, multiscale-entropy *shape* parameters (decay rate under cognitive challenge) are known to carry long-lived developmental-programming signatures in ECG-only recordings. Third, perturbation response profiles (cold pressor, orthostatic tilt, TSST) probe the *curvature* of the action landscape, not merely its minimum, and should reveal regime-dependent shifts in the empirical-coupling product that resting-state metrics miss.

## 5 Limitations

This is a reanalysis of a two-animal pilot cohort (Castel et al., 2024), with one VNS-treated and one LPS-only animal. With *N* = 2 no inferential between-arm statistics are possible; all reported physiological observations are case-series descriptions and must be regarded as hypothesis-generating. The pre-specified null comparisons in §2.6 (Poisson, bootstrap, Gaussian-CV, parameter sensitivity, phase-randomised VENG surrogates, *R* versus simpler ICI-series baselines, and direct VENG–RR phase locking) test whether the central derived metrics carry signal beyond their most obvious null alternatives. They are not substitutes for between-animal replication. Two of the seven companion tests returned negative or borderline results that we report explicitly: (a) the changepoint detector exhibits a sharp threshold cliff near the default value of 2.05 (*n*_CP_ halving with a +15% change), motivating per-animal threshold calibration in the prospective cohort rather than a fixed value. Because every downstream coordination-dynamics quantity in this paper—changepoint count, ICI distribution, *R*, HKB *k*, Arnold-tongue position, CSD indicators, CAP even/odd assignment, and the *R* · eScalE product—is conditional on the detected changepoint sequence, the threshold cliff implies that the present physiological trajectories should be read as conditional on the stated detector operating point rather than as parameter-free properties of the raw VENG signal. We have not performed a comprehensive trajectory-stability analysis across thresholds because re-running the detector with the present implementation is computationally expensive (one parameter sweep on two recordings took ∼45 minutes on a single core); a faster implementation suitable for cohort-wide robustness sweeps is on the methods companion’s roadmap, and (b) the direct phase-locking test of the LF/HF bridge (§3.8) returned a null at all 16 testable timepoints, indicating that the present 5-min recording duration is inadequate for that test even if the hypothesis is correct. The absolute changepoint counts depend on detector parameters (bandwidth, threshold) and should not be compared across studies without reference to the full parameter set, which we release. The parameter-sensitivity sweep reported in §2.6 (Fig. 3) covers two representative recordings; a cohort-wide sweep at every timepoint is on the methods-companion roadmap but was not feasible at the present implementation’s runtime cost. Piglet 3 recordings lack physical calibration and are analysed from raw digital samples; within-animal temporal dynamics are valid but absolute VENG amplitude is not. Piglet 3 left has no baseline; Piglet 3 right is missing the 120-min timepoint. The *R* · eScalE regime-dependent coupling is observed in *N* = 1 VNS-treated animal (Piglet 3 ECG quality precludes the comparison) and must be regarded as a post-hoc empirical observation; the protocol in §6 is designed specifically to falsify it. The CIMVA metrics in the current ECG analysis are limited by the signal quality of the piglet cohort’s ECG channels and should be reapplied to longer, higher-quality recordings as part of the prospective validation. Specific physiological labels for the two latent states (e.g. afferent vs. efferent phases of a vagal duty cycle) are interpretive proposals discussed in §4 and are not established by the present data; the data-supported claim is the narrower one that the two states are functionally distinct at the level of high-frequency CAP band power.

## 6 Proposed Prospective Validation

We propose a pre-registered five-phase protocol for the next cohort (*n* ≥ 8 per arm, with high-quality ECG recording ensured throughout): (i) 60-min bilateral VENG + ECG baseline to establish per-animal *R*, HKB parameters, and baseline *R* · eScalE; (ii) first LPS exposure with continuous recording through acute resolution; (iii) 24–72 h recovery until standard HRV metrics normalise; (iv) post-recovery re-assessment of HKB potential, Arnold-tongue position, and *R* · eScalE, testing whether the moderate-regime product (≈0.33 in Piglet 2) and the VNS-overshoot product (≈0.80) replicate across animals; (v) a second, lower-amplitude challenge (LPS or cold pressor) to probe response dynamics.

### Primary prediction (biphasic VNS-response trajectory)

VNS-treated animals will show an acute reduction in changepoint count at the first post-LPS timepoint relative to LPS-only animals, followed by recovery and overshoot exceeding baseline; the LPS-only and LPS + VNS trajectories will cross at approximately 60–75 min post-LPS. This prediction is directly motivated by the within-cohort observation that Piglet 2 sits below Piglet 3 right for the first 60 min post-LPS (§3.3) and is the strongest available falsifier of the biphasic-divergence reading: an alternative model in which VNS protects switching at every timepoint would predict no crossover, while a null model would predict no consistent ordering at any timepoint.

### Secondary prediction (regime-dependent *R* · eScalE coupling)

The two-regime structure— moderate-regime trade-off with stable *R* · eScalE, interrupted by VNS-overshoot doubling— will replicate across VNS-treated animals, with the magnitude of the overshoot scaling with VNS dosage parameters.

### Tertiary prediction (CSD early-warning)

Critical-slowing-down indicators applied to the second challenge will reveal reduced resilience (shallower HKB barrier Δ*V*) in previously exposed animals, and the same indicators applied to RR-interval series should flag the bifurcation in advance through the LF/HF bridge proposed in §3.8. The latter requires direct phase-locking analysis between simultaneous VENG and RR series in the prospective cohort and is the primary methodological test of the LF/HF-bridge hypothesis.

### Critical quality control

The prospective cohort must ensure ≥200 clean beats per 5-min window (the current Piglet 3 cohort failed this criterion at all but one timepoint, rendering the empirical-coupling test inconclusive for the untreated arm) and per-animal pre-LPS baselines bilaterally to permit within-animal laterality analysis.

## 7 Conclusion

We have presented a modular open-source Python pipeline that extracts coordination-dynamics descriptors—spectral-KL changepoints, phase-coherence index *R*, HKB bistable-potential parameters, Arnold-tongue position, critical-slowing-down indicators, and a joint VENG–HRV empirical-coupling scan—from simultaneously recorded vagus nerve electroneurogram and ECG, with pre-specified null comparisons that test whether the central derived metrics carry information beyond their most obvious null alternatives. Applied to the two-animal neonatal piglet endotoxaemia cohort of Castel et al. (2024), the pipeline recovers the preparation’s known features and surfaces three case-series observations: (i) qualitatively divergent coordination-dynamics trajectories under LPS versus LPS + VNS that cross at 60–75 min post-LPS; (ii) a >100-fold pre-collapse rise in inter-changepoint-interval variance that exceeds bootstrap nulls drawn from the same animal’s pre-collapse pool; and (iii) a regime-dependent empirical coupling between VENG coherence and HRV complexity in the VNS-treated animal (*R* · eScalE ≈ 0.33 during moderate challenge, ≈0.80 during VNS-overshoot). We frame all three as hypothesis-generating observations and not inferentially established findings; the manuscript closes with a pre-registered prospective protocol designed specifically to falsify them and connect them to the broader vertical-organizing-principles framework recently proposed in this Journal (Frasch, 2026).

## Data and code availability

Full Python pipeline: github.com/martinfrasch/veng_minaction (currently private during peer review; access on request, public release planned on acceptance under MIT licence). All parameters in §2 are the defaults shipped with the repository. Raw recordings analysed here are those of Castel et al. (2024) (preprint Castel et al. 2020) and are available from the corresponding author of that dataset on reasonable request, subject to institutional data-use agreements.

## Acknowledgments

We thank the full experimental team of the neonatal piglet sepsis cohort whose protocol and dataset (Castel et al., 2020, 2024) made this reanalysis possible. No new *in-vivo* experiments were performed for this study.

## Author contributions

M.G.F. conceived the pipeline, ported the changepoint detector to Python, implemented the coordination-dynamics, CAP, and empirical-coupling modules, performed all analyses, and drafted the manuscript. M.L. provided the original R changepoint code base and verified numerical equivalence of the Python port against the R progenitor. P.B., G.F., and A.D. designed and performed the original neonatal piglet cohort experiment whose data is reanalysed here (see Castel et al. 2024) and advised on interpretation of the physiological results. All authors read, revised, and approved the final manuscript and agree to be accountable for all aspects of the work.

## Preprint statement

This manuscript has been deposited on bioRxiv as a preprint (DOI to be assigned) and submitted to *The Journal of Physiology* via the bioRxiv direct-transfer option.

